# Green Synthesis of a Novel Gelatin Crosslinked Oxidized Pullulan-Lignin biocomposite film for active food packaging

**DOI:** 10.1101/2025.05.06.652438

**Authors:** Ayan Banerjee, Mohit Kumar Mehra, Althuri Avanthi

## Abstract

Biopolymer-based plastics, made from plants and microbes, are gaining popularity in food packaging due to their eco-friendliness, but degrade upon contact with water, thus requiring the addition of water-resistant materials. Lignin was extracted from cotton stalk using the acid-alkali method and was characterized through FTIR and 2D HSQC NMR to evaluate its structure and purity. Pullulan dialdehyde, gelatin, and lignin were blended using a green process to make water-stable PGL hydrogel film. Incorporation of Lignin into PDA/Gelatin blend, having imine bonds, increased network strength and solvent-resistance by extensive physical interactions like H-bonding and electrostatic interactions, which were confirmed through FTIR and XPS analysis. These films were cured at 60°C, making the process energy efficient. PGL film exhibited water stability, high mechanical strength, moisture barrier, and bioactive properties ideal for fruit preservation. Lignin improved hydrophobicity, hydrogel strength, biodegradability, and non-flammability relative to the control. The films’ effectiveness for food packaging was assessed by analyzing the impact on the preservation and shelf life of kinnow oranges. Physical properties, such as weight loss, firmness, and chemical properties, including pH, titratable acidity, lipid oxidation, sucrose content, antioxidant activity, and ethylene concentration of oranges before and after packaging, were monitored. PGL film showed high stability in hot and ambient water, with a water swelling index of 303%, impressive tensile strength of 8.5 MPa, and elongation at break exceeding 72%. PGL-packed oranges produced 0.034±0.0032 ppm/g of ethylene after seven days, which is 4.2-fold lower than unpacked oranges, signifying the fruit preservation ability of PGL films.

## 1. Introduction

Conventional plastic packaging materials, including polyethylene (PE), polypropylene (PP), polyvinyl chloride (PVC), and polyethylene terephthalate (PET), are procured primarily from petroleum and natural gas, which are non-renewable in nature (Bauer et al., 2022). The high dependence on these resources contributes to environmental concerns due to greenhouse gas emissions, non-biodegradability, and plastic waste accumulation (Bauer et al., 2022). Furthermore, synthetic plastics merely provide moisture resistance, food containment, and easy transport but lack active functionalities (Shiva et al., 2024). The passive nature of these synthetic plastics necessitates the addition of chemical preservatives for food preservation (Shiva et al., 2024) (Rodrigues et al., 2022). However, active packaging is needed to extend food shelf life, which can keep food fresh and healthy and reduce the need for preservatives by providing properties like antimicrobial, antioxidant, and UV-blocking effects.

To address these limitations, developing biodegradable and active food packaging materials has gained significant attention. Various biopolymer-based films, including polysaccharides (e.g., starch (Yuan et al., 2025), pullulan (Trinetta & Cutter, 2025)), proteins (Sun et al., 2025), and polyhydroxyalkanoates (PHA) (Amir et al., 2025), have been explored for their sustainability and bioactive properties. However, challenges such as inadequate mechanical strength, hydrophilicity, and limited moisture barrier properties hinder their widespread application (Taherimehr et al., 2021).

Pullulan, an exopolysaccharide produced by *Aureobasidium pullulans*, exhibits excellent film-forming ability and oxygen barrier properties but suffers from poor water resistance and low mechanical strength (Ghosh et al., 2022) (Farris et al., 2014). Cotton is a popular plant worldwide for its application in the textile industry (Cai et al., 2024). Apart from its use in textiles, it is also a source of edible vegetable oils (Ganesan et al., 2018), which provide human nutrition. Currently, cotton is planted in over 80 countries, including India, Australia, USA, Egypt, and Brazil (Jindal et al., 2023). Cotton stalk is an abundant agricultural residue, which is produced about 30 million tonnes per annum in India. (Jindal et al., 2023). Conventional on-field practice to alleviate cotton stalk after harvest is to burn in open, increasing its carbon footprint. CS has 21-25% lignin content (G. Li et al., 2022) making it a potential lignin source.

Lignin is an aromatic second-most-abundant, and heterogenous biopolymer with a polyphenolic structure (Garg & Avanthi, 2025) (Rico-García et al., 2020).

Lignin is mainly used as a filler or low-grade additive to boiler fuel for energy production (). It consists of subunits like guaiacyl (G), syringyl (S), and p-hydroxyphenyl (H) (Halloub et al., 2022), whose composition changes with the change in biomass type (Avanthi & Banerjee, 2016). The heterostructure of lignin makes it challenging to develop lignin-based products (Du et al., 2022). Despite its complex structure, it shows properties like antioxidant, anti-microbial, hydrophobicity, and flame-retardant properties. It possesses high mechanical strength due to its inner crosslinked structure, and also photothermal capabilities(Liu & Bernaerts, 2024).

However, unstable H-bonding between lignin and pullulan limits the film’s stability in aqueous environments. Thus, a crosslinker or copolymer, which will bind pullulan and lignin together, is needed. Toxic crosslinkers like formaldehyde-based (glutaraldehyde (Liu & Bernaerts, 2024)), epichlorohydrin (Dessouki et al., 2001) and isocyanates (Shibata et al., 2001) are not suitable in synthesizing food packaging films as they can leach into food items (Mugnaini et al., 2023). Several green crosslinkers, like citric acid (Wen et al., 2021), tannic acid (Zhao et al., 2025), and cyclic phosphates (Dulong et al., 2011), are being used to fabricate different food packaging films yet have not been reported to impart water stability to the films. Various copolymers, like PVA (Min et al., 2021), cellulose (Min et al., 2021), starch (M. Zhang et al., 2023), and Chitosan (M. Zhang et al., 2023), have also been used to enhance physical interactions like hydrogen and electrostatic bonding.

Gelatin is a widely available, low-cost protein that is derived from animal collagen (Gasti et al., 2022). It is classified into Type A Gelatin (acid hydrolysis) and Type B Gelatin (alkaline hydrolysis) (Xiao et al., 2021). Type A Gelatin has a higher protein content contributing to UV-C light absorption ability. However, Gelatin (Type A and Type B) exhibit poor moisture resistance, rendering the film highly unstable under high moisture environment. Type A has lower viscosity, and better ability to form films than Type B. Hydrogen bonding between carbonyl and amine groups results in triple-helix structure of gelatin that exhibits high mechanical strength (Gómez-Guillén et al., 2011).

Although lignin reacts electrostatically with gelatin by disrupting the stable intermolecular H-bonding in the 3D protein structure of gelatin (Aadil et al., 2016), the lignin/gelatin blend is partially water-stable. On the other hand, only Pullulan and gelatin also do not form stable films due to lack of chemical interactions. It is reported that Pullulan dialdehyde (PDA) readily reacts with gelatin to form stable covalent imine bonds (L. Zhang et al., 2019). Oxidation of Pullulan increases the reaction sites by introducing aldehyde groups, that readily reacts with other copolymers and increase water stability and antibacterial properties (Roy et al., 2023).

This study developed a novel PGL biocomposite film composed of Pullulan dialdehyde (P), gelatin (G), and lignin (L) (Fig. 1), as an eco-friendly alternative to synthetic plastic packaging. Lignin was extracted from cotton stalks using an acid-alkali method, while pullulan was oxidized to pullulan dialdehyde (PDA) to enhance crosslinking efficiency. Gelatin was incorporated as a biodegradable crosslinker to improve film stability and mechanical strength. The film fabrication method follows all 12 principles of green chemistry using green, nontoxic solvents and sustainable chemistry. This novel PGL films’ detailed structure (Fig. 2A), properties like water stability, moisture barrier, mechanical, and rheological properties were thoroughly evaluated to assess their efficacy in fruit packaging. The fabricated PGL films were tested for active packaging of citrus fruit variety with short shelf life. The nutrient contents, chemical, and physical properties of kinnow oranges change drastically post-harvest due to their limited shelf life compared to regular oranges (Challana et al., 2025). Preservation during storage and transportation of these varieties is highly challenging, which is attempted to be addressed in this study. Ethylene gas, which is a biomarker of fruit ripening, was analyzed to evaluate the fruit preservation efficacy of PGL film.

**Fig. 1.**
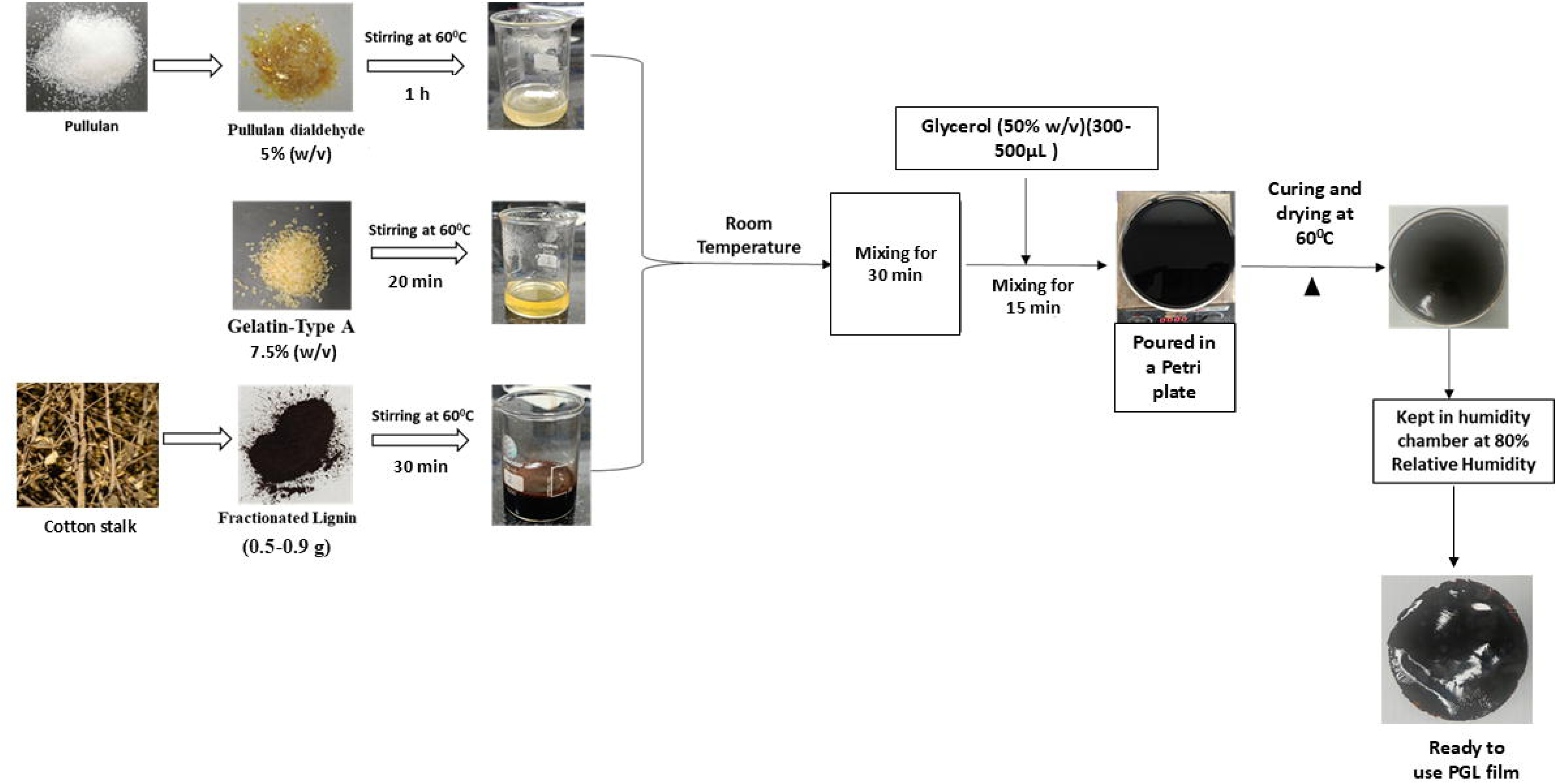
Schematic representation of PGL film preparation.

**Fig. 2.**
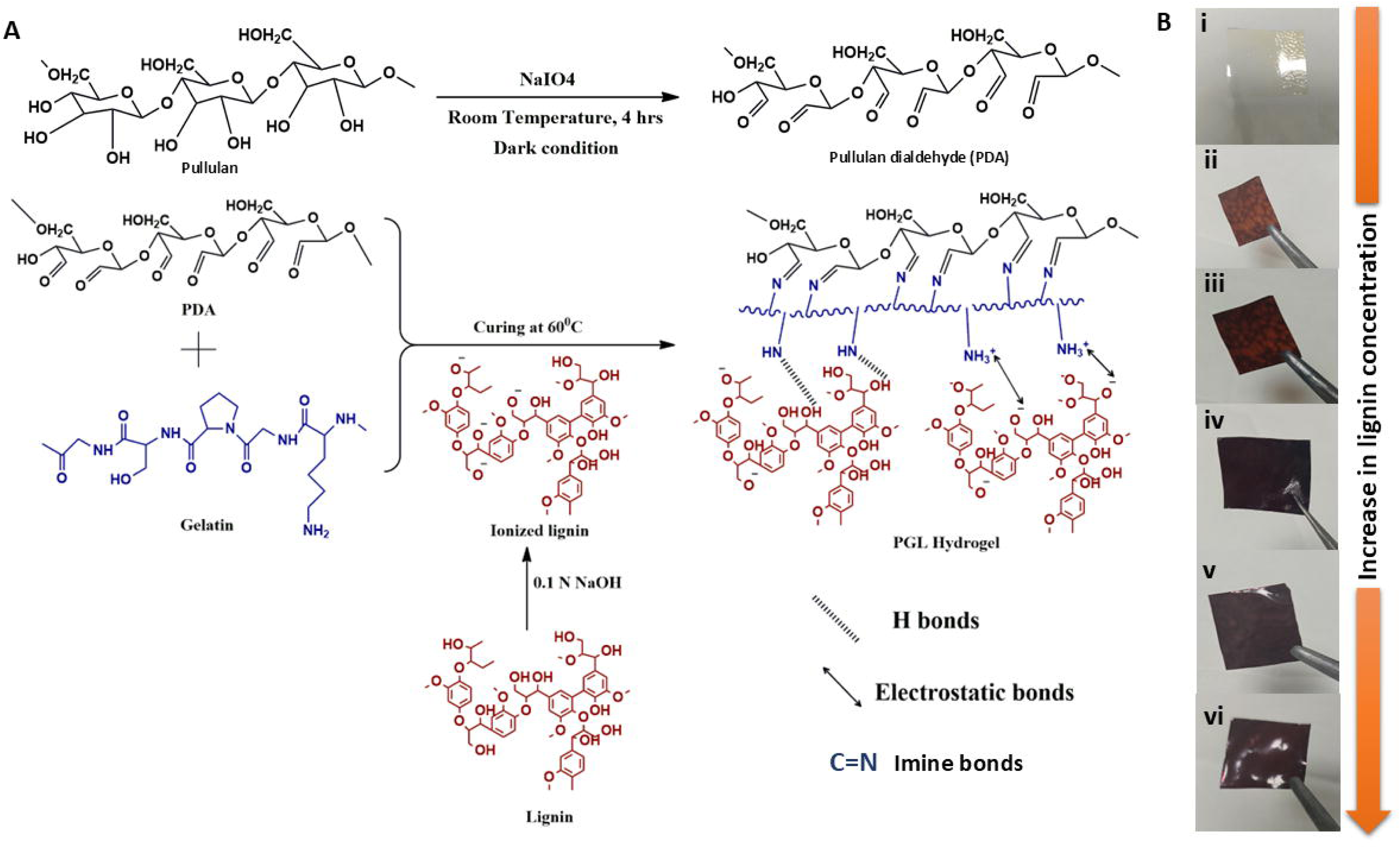
(A) Reaction mechanism of the formation of PGL hybrid biocomposite, showing imine bonds, hydrogen bonding, and electrostatic interactions; (B) Pictures of (i) control film and films with varying Lignin amounts (ii) 0.5g, (iii) 0.6g, (iv) 0.7g, (v) 0.8g, (vi) 0.9g.

The biodegradation ability of PGL film in both soil and water was evaluated to ensure environmental sustainability.

The resulting PGL films can exhibit enhanced water resistance, antimicrobial activity, UV shielding, and radical scavenging properties, making them suitable for active fruit packaging. This green and eco-friendly film is novel, and there are no studies in the literature with gelatin crosslinked hybrid pullulan-lignin biocomposite to the best of our knowledge.

## 2. Experimental section

### 2.1. Materials

Cotton stalk (CS) was collected from local farmers of Sangareddy, India (latitude and longitude 17.602502517051256, 78.04290842650505). Sodium Hydroxide (NaOH), Sulfuric acid (H_2_SO_4_), Sodium metaperiodate (NaIO_4_), Glycerol, DPPH (2,2-diphenyl-1-picrylhydrazyl), Nutrient Broth, Agar, de Man, Rogosa, and Sharpe (MRS) broth, phenol, Trichloroacetic acid (TCA) and Thiobarbituric acid (TBA) were supplied by Sisco Research Laboratories Pvt. Ltd. (SRL), India. Ethylene glycol and hydroxylamine hydrochloride were bought from ChemSynth Fine Chemicals, India. Dialysis membrane (MWCO: 12,000–14,000 Da) and Gelatin Type A were procured from HiMedia Laboratories Pvt. Ltd., India. Pullulan was obtained from TCI Chemicals, India. Kinnow Oranges were bought from a local shop of the Indian Institute of Technology, Hyderabad (latitude and longitude 17.585286885207577, 78.11976144726191).

### 2.2. Lignin extraction from agricultural waste

Compositional analyses of the CS, including % extractives, % cellulose, % lignin, and % ash, was done by following NREL protocol (Supporting information Table S1) (Althuri et al., 2017), (Sluiter et al., 2005) (Sluiter et al., 2008). Lignin was extracted by the acid-alkali method under different reaction conditions. For this, 10 g of powdered CS was taken, having a particle size of ≤ 0.25 mm. Hemicellulose was removed by acid pretreatment with 1-3% H_2_SO_4_ (v/v) at a temperature of 100-120°C for 15-120 min in an autoclave. After acid pretreatment, the residue was filtered and neutralized, followed by drying in an oven at 50-60 °C overnight. The dried residue was treated with 3-5% NaOH (w/v) at 100-120 °C for 15-120 min. Each experimental run had solid loading (% w/v) between 5-10 %. After NaOH treatment, vacuum filtration was done to separate the black lignin liquid from the residue. Concentrated H_2_SO_4_ was added dropwise to the alkaline lignin solution to precipitate lignin. The precipitated lignin was washed 3-4 times with distilled water until neutral pH, and dried to obtain fractionated lignin at 60°C overnight. The purity of extracted lignin was evaluated following the NREL protocol.

### 2.3. Enhancing lignin yield and purity by statistical analysis

The Minitab 17 software was used to create a Plackett-Burman (PB) experimental design, which explores the key factors influencing lignin yield from cotton stalk using the acid-alkali method. A total of 12 experimental runs were carried out (Table 1), each triplicated, encompassing various combinations of factors at two levels: low (−) and high (+). These factors included solid loading (5%, 10 % w/v), sulfuric acid (H_2_SO_4_) concentration (1%,3% v/v), Acid pretreatment time (15 min,120 min), Alkali percentage (3%,5% w/v), and Alkali treatment time (15min, 120min). After completing each experimental run in triplicates, statistical analysis was done using the ANOVA, Half Normal plot, and Pareto Chart to estimate the significance of the overall model and each factor on lignin yield and recovery.

**Table 1.**
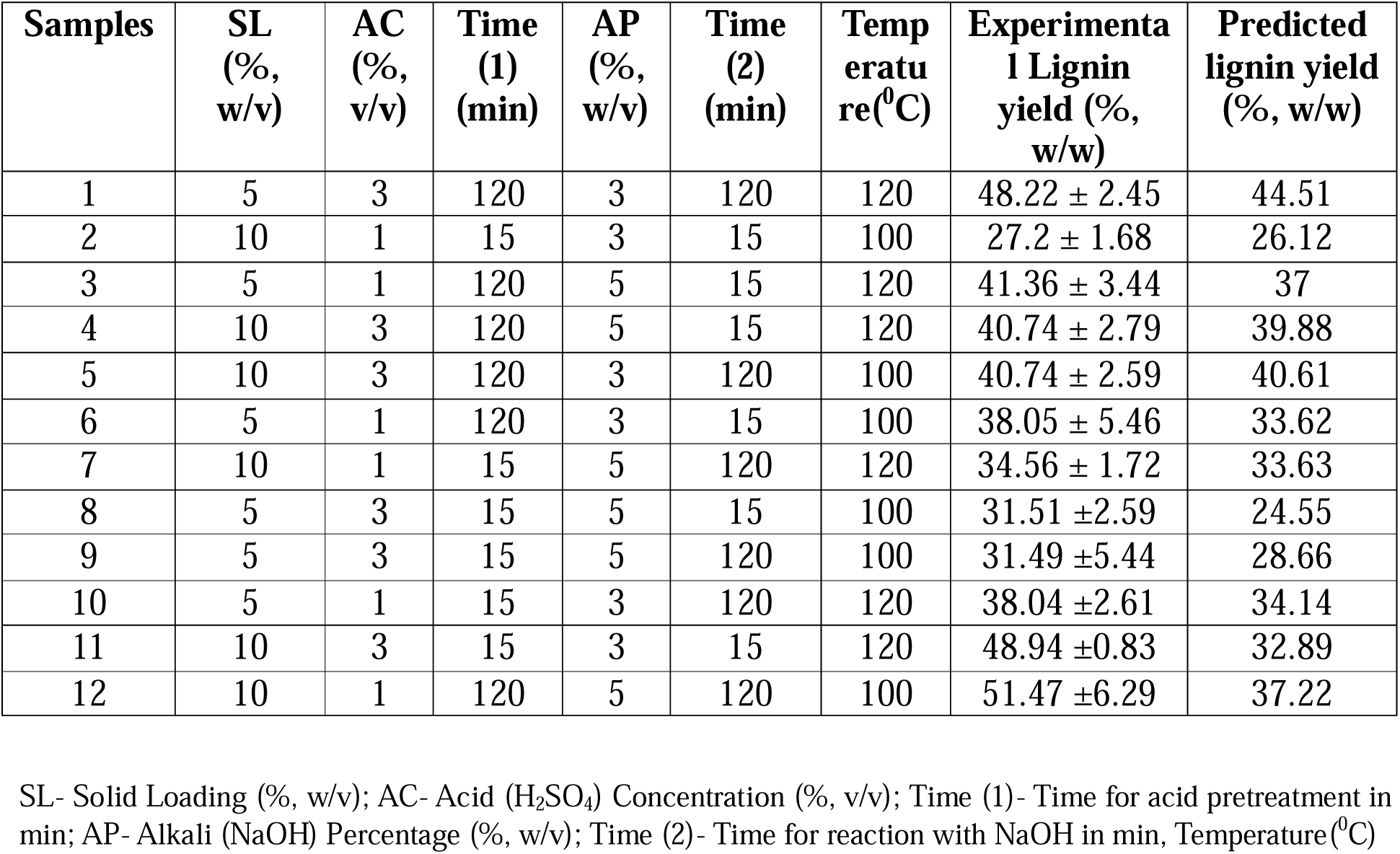
Plackett-Burman (PB) design for improved Lignin extraction from cotton stalk by acid-alkali method.

| Samples | SL<br>(%,<br>w/v) | AC<br>(%,<br>v/v) | Time<br>(1)<br>(min) | AP<br>(%,<br>w/v) | Time<br>(2)<br>(min) | Temp<br>eratu<br>re(<sup>0</sup>C) | Experimenta<br>l Lignin<br>yield (%,<br>w/w) | Predicted<br>lignin yield<br>(%, w/w) |
| --- | --- | --- | --- | --- | --- | --- | --- | --- |
| 1 | 5 | 3 | 120 | 3 | 120 | 120 | 48.22 ± 2.45 | 44.51 |
| 2 | 10 | 1 | 15 | 3 | 15 | 100 | 27.2 ± 1.68 | 26.12 |
| 3 | 5 | 1 | 120 | 5 | 15 | 120 | 41.36 ± 3.44 | 37 |
| 4 | 10 | 3 | 120 | 5 | 15 | 120 | 40.74 ± 2.79 | 39.88 |
| 5 | 10 | 3 | 120 | 3 | 120 | 100 | 40.74 ± 2.59 | 40.61 |
| 6 | 5 | 1 | 120 | 3 | 15 | 100 | 38.05 ± 5.46 | 33.62 |
| 7 | 10 | 1 | 15 | 5 | 120 | 120 | 34.56 ± 1.72 | 33.63 |
| 8 | 5 | 3 | 15 | 5 | 15 | 100 | 31.51 ± 2.59 | 24.55 |
| 9 | 5 | 3 | 15 | 5 | 120 | 100 | 31.49 ± 5.44 | 28.66 |
| 10 | 5 | 1 | 15 | 3 | 120 | 120 | 38.04 ± 2.61 | 34.14 |
| 11 | 10 | 3 | 15 | 3 | 15 | 120 | 48.94 ± 0.83 | 32.89 |
| 12 | 10 | 1 | 120 | 5 | 120 | 100 | 51.47 ± 6.29 | 37.22 |

SL- Solid Loading (% , w/v); AC- Acid (H<sub>2</sub>SO<sub>4</sub>) Concentration (% , v/v); Time (1)- Time for acid pretreatment in min; AP- Alkali (NaOH) Percentage (% , w/v); Time (2)- Time for reaction with NaOH in min, Temperature(<sup>0</sup>C)

### 2.4. Pullulan modification to pullulan dialdehyde

Pullulan dialdehyde (PDA) was synthesized by slightly modifying the method reported by Wang et al. (H. Wang et al., 2016). Pullulan aqueous solution (2% (w/v)) was prepared in deionized water. Sodium metaperiodate (1.3% (w/v)) was added to the aq. Pullulan solution, and the reaction mixture was stirred continuously in the dark at room temperature for 4 h. To terminate the reaction, 10 mL of ethylene glycol was added. The solution was dialyzed against deionized water for 24 h using a dialysis membrane (MWCO: 12,000–14,000 Da) to remove unreacted sodium metaperiodate. The purified fraction was dried in a hot air oven at 60 °C overnight. The presence of aldehyde groups in the oxidized pullulan was confirmed by FTIR spectroscopy.

### 2.5. Degree of oxidation and Aldehyde content of PDA

#### 2.5.1. Degree of oxidation

The degree of oxidation of pullulan was determined using UV/Vis spectroscopy at 222 nm (Simon et al., 2022). The concentration of unreacted periodate was measured by taking 1 mL of post-dialysis filtrate. The absorbance values were kept within the range of 0.5 to 1.1 by adjusting the dilution factor. Jenway UV/Vis spectrophotometer was used for these measurements with distilled water as a blank. The DO% was calculated using the following equation (Eq. 1), assuming no side reactions.

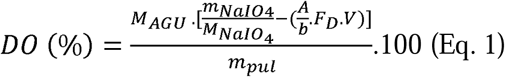

where *m_NaIO_*_4_ and *m_pul_* are the mass of sodium periodate and pullulan, respectively; *M_NaIO_*_4_ is the molecular weight of the sodium periodate, *A* the arithmetic means of the measured absorbance, *b* the calibration curve slope, *F_D_ is* the dilution factor, and *V is* the solvent (deionized water) volume; *M_AGU_* is the molecular weight of the anhydroglucose unit, i.e. 162.14 g/mol.

#### 2.5.2. Aldehyde Content

The molar content of aldehyde present per g of PDA was determined by using titration method (V. Kumar & Yang, 2002). Hydroxylamine hydrochloride reacted with aldehyde groups of PDA to form Schiff base and hydrochloric acid. In this process, 300 mg PDA was initially dissolved in 25 mL distilled water, and the solution was mixed with 20 mL of 1.5 % (w/v) hydroxylamine hydrochloride at 40°C for 4 hrs at pH 5. The pH of the mixture was checked and titrated against 0.1 mol/L NaOH aq. solution. The volume of hydrochloric acid neutralized by NaOH solution in the sample is recorded as V_c_. Similarly, the volume of NaOH solution required for the pure pullulan (unmodified) under the same conditions was recorded as V_b_, which was used as a blank control. Therefore, the aldehyde content and percentage of PDA were calculated using Eq. 2& 3:

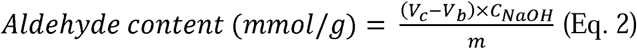

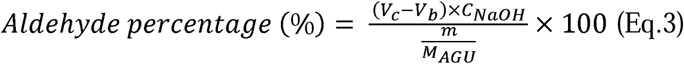

Where C_NaOH_ = NaOH concentration (0.1 mol/L), m= oven-dried mass of oxidized pullulan (0.3g), and *M_AGU_* is the molecular weight of the anhydroglucose unit, i.e., 162.14 g/mol.

### 2.6. Synthesis and fabrication of novel PGL biocomposite

1 g of PDA was dissolved in 20 mL of distilled water at 60°C for 1 hour to form a uniform solution. Lignin at different concentrations (0.5 g-0.9 g, 20-40% w/w) was dissolved separately in 20 mL NaOH solution (0.1 N) until fully dissolved. Meanwhile, 1.5 g of gelatin was mixed with 20 mL of distilled water at 60°C until dissolved and cooled to room temperature. PDA solution (20 mL) was gradually added to the gelatin solution under stirring and mixed for 30 min. Lignin solution (20 mL) was added drop by drop to the PDA-gelatin blend and stirred at room temperature on a mechanical stirrer for another 30 min. Glycerol (50% w/v) was used as plasticizer at different volumes (300-500 μL) was added to the PGL solution and stirred for 10 minutes. PGL solution was poured into a 15 cm diameter petri dish, which was placed on a hot plate set at 60°C for curing and drying. After 1 h, once the crosslinked hydrogel dried, the petri dish was transferred to a humid chamber with 80% relative humidity and left overnight (Fig. 1). The next day, the dried films were carefully peeled off and stored under the same humid conditions for 3 days to prepare them for further testing and characterization.

PGL films, represented by Pullulan dialdehyde (P), Gelatin (G), and Lignin (L), were set to range within a ratio of 2:3:1 to 10:15:9 (w/w) (Fig. 2B(ii-vi)). The specific PGL formulations include 2:3:1 for PGL-0.5, 10:15:6 for PGL-0.6, 10:15:7 for PGL-0.7, 10:15:8 for PGL-0.8, and 10:15:9 for PGL-0.9 films, respectively. PDA/gelatin film was used as a control (Fig. 2B (i))

**Fig. 3.**
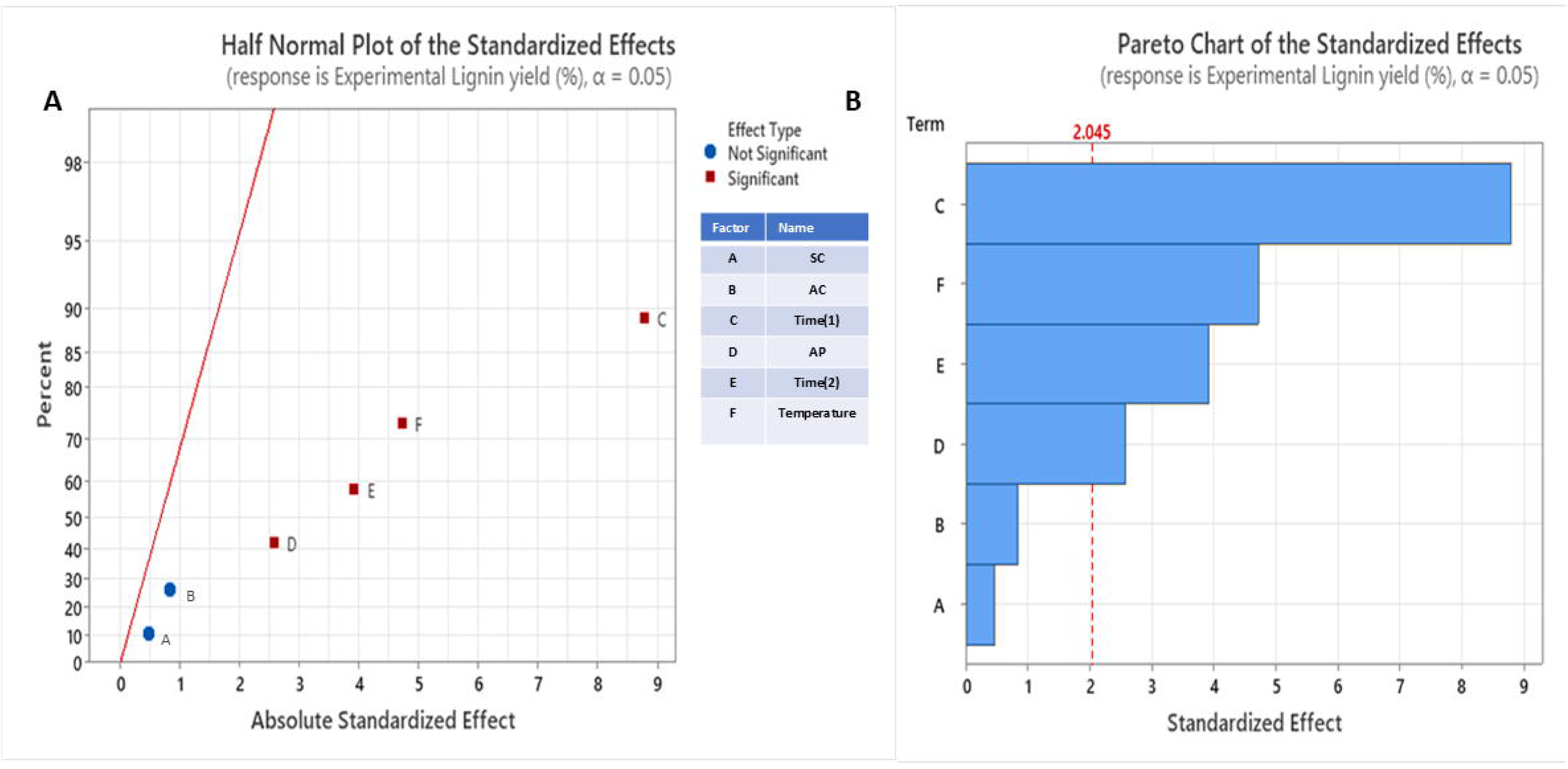
Half Normal plot and Pareto Chart of the standardized effects of lignin extraction by acid-alkali process.

### 2.7. Chemical Characterizations

#### 2.7.1. 2D Heteronuclear Single Quantum Coherence Nuclear Magnetic Resonance (HSQC NMR)

Bruker Ultra Shield 400 spectrometer was used to record the 2D HSQC spectra of fractionated lignin. In 0.6 mL of DMSO-d6 (99.8% deuterated) 35 mg of lignin was dissolved. For quantitative analysis, the standard Bruker pulse software hsqcetgp was utilized. For the ¹H dimension, the spectral range was set to 0 to 8 ppm, and for the ¹³C dimension, it was set to 0 to 180 ppm. With a one-second recycle delay, the ¹H dimension was scanned thirty-two times. There were 3072 transients in the ¹³C dimension with a 2-second relaxation delay. Coupling constant (¹JCH) of 145 Hz was used. The data matrix was zero-filled to 1024 points in the ¹³C dimension prior to the Fourier processing. MestReNova-15.0.1 software was used to process the spectral data.

#### 2.7.2. Fourier Transform Infrared Spectroscopy (FT-IR)

Chemical characterization of PDA, fractionated (extracted) lignin, control (PDA/gelatin film), and PGL films was done by using FT-IR spectroscopy using a Bruker TENSOR 37 FTIR spectrophotometer. The spectra were recorded from 400 to 4000 cmC¹(Garg & Avanthi, 2025).

#### 2.7.3. X-ray photoelectron spectroscopy (XPS)

To explore more insights regarding the interactions in both the control and PGL films, AXIS Supra-Kratos analytical spectrophotometer coupled with a monochromatic Al K_α_ X-ray source (1486.6 eV) was used to perform the XPS analysis. The experiments were conducted utilizing 1 mm spot size and a detection depth of 5 nm. High-resolution spectra were acquired via an energy analyzer with a pass energy of 40 eV. The binding energy scale was calibrated using the C1 peak at 284.6 eV. Peak fitting and data analysis were performed using Origin software.

### 2.8. Water swelling property

The water swelling property assesses the films’ long-term water stability as well as resistance to water absorption. Conditioned films (PGL-0.5-0.9 and control films) were cut into 2 × 2 cm² pieces, and the initial dry mass (W_d_) of each was noted. Water swelling ratio was recorded over 1000 min by immersing the samples in 25 mL of distilled water (L. Wu et al., 2019). Whatman filter paper was used to remove excess water from the film samples, which were removed at specific intervals, and the swollen weight (W_s_) was recorded. Using Eq.4 water swelling ratio was calculated.

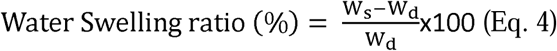

### 2.9. Water Vapor Transmission Rate (WVTR)

Water Vapor Transmission Rate (WVTR), which signifies the moisture barrier property of the films, was measured (Garg & Avanthi, 2025). Films were cut into 3 x 3 cm² pieces and fitted over the opening of a 150 mL conical flask containing 10 mL distilled water and secured with Parafilm, and its initial weight (W_1_) was noted. After an incubation time of 24 h at 40 ± 2°C, the flask was taken out, cooled down, and weighed to determine the final weight (W_2_).

The WVTR (g/m²·h) was calculated using the following Eq. 5, where T represents the time of incubation and A specifies flask opening area (cm²).

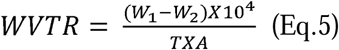

### 2.10. Mechanical testing

Using an Instron mechanical testing machine, the stress-strain response of the control and PGL films was studied according to ASTM D882, a standard for films thinner than 1 mm. The films were cut into rectangular samples measuring 50 mm × 25 mm for testing, and they were attached between the machine grips. The crosshead speed was maintained at 10 mm/min, and the initial grip distance was 10 mm. Micrometer was used to measure each film’s thickness. Tensile strength (MPa) and elongation at break percentage (%) were calculated from the stress-strain curves. Toughness, which is a measure of material’s ability to absorb energy before fracture, was calculated from under the curve of stress-strain plot.

### 2.11. Bioactive properties

#### 2.11.1. UV-Blocking property

Lambda 365 UV-Vis spectrophotometer was used to analyse the UV blocking property of films. Film samples were cut into 2 × 2 cm² pieces, with air as the reference, for the analysis. The absorbance and transmittance values of control and PGL film samples were recorded between 200 to 800 nm range (Garg & Avanthi, 2025). Sun Protection Factor (SPF) (Almeida et al., 2019) was determined in the most harmful UV region (UV-B) by the following Eq. 6. From the SPF value the UV-B blocking percentage was also calculated to quantify the films’ UV-blocking efficiency using Eq. 7

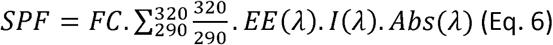

Where FC is a constant value of 10, EE represents the erythemogenic effect, I denotes the intensity of sunlight, and Abs corresponds to the absorbance of the sample. Absorbance measurements were taken within the 290–320 nm at 5 nm intervals and applied to Eq. 6, the constant EE (λ).I (λ) with respect to wavelength (290-320 nm), predefined by Mansur (1984) (Almeida et al., 2019), was used for calculating SPF.

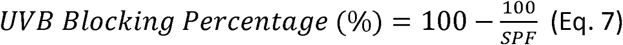

Further, the UVA/UVB ratio of the films was calculated (You et al., 2023) by integrating the area under the curve to check the broad-spectrum UV protection capacity.

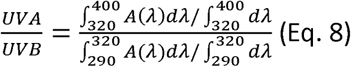

#### 2.11.2. Radical Quenching Effect

To assess the antioxidant or radical scavenging / quenching properties of PGL and control films, 200 mg of film sample was added to 10 mL of 95% ethanol and allowed to stand at 50°C for 3 h in the dark. After incubation, 1.5 mL of 0.06 mM DPPH (2,2-diphenyl-1-picrylhydrazyl) in 95% ethanol was mixed with 1.5 mL of the supernatant. The mixture was agitated at room temperature without light for 30 min. Jenway UV/Vis spectrophotometer was used to test the solution’s absorbance at 517 nm. Similarly, a blank sample was made, without the film. Eq. 9 was used to determine the radical scavenging activity (%), and the experiments were carried out in triplicate (Garg & Avanthi, 2025)

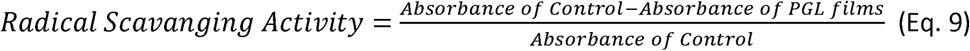

#### 2.11.3. Antimicrobial activity

Disk diffusion method (Garg & Avanthi, 2025) was employed to evaluate the antimicrobial efficacy of film samples against *Lactobacillus pentosus, Lactobacillus rhamnosus, and Lactobacillus casei*. To ensure sterility, films with diameters of 0.5 cm (PGL-0.7 film) and 0.6 cm (Control film) were first subjected to UV radiation for 15 minutes. The samples were then placed on agar plates with 10 µL of MRS broth inoculum of test microbial strains.

### 2.12. Surface morphology of films

The film samples were coated with gold by a sputter instrument. The gold deposition time was 30s. Gold-coated samples were mounted on carbon tapes and examined in a JEOL JIB 4700F FIB-SEM at 3-5 kV. The elemental studies were also done by an inbuilt EDX to check further film impurities.

### 2.13. Surface wettability

Contact angle of dried test and control films were measured to check surface wettability using a contact angle meter (KRUSS DSA-25S). Water droplet (4 μL) was deposited on the surface of films, and the water contact angle was measured after few seconds. These experiments were done in triplicates.

### 2.14. Rheological studies

Rheology of film-forming solution (PGL-0.7 and control) was assessed using a Kinexus Pro+ (Malvern Panalytical, UK) rheometer using parallel discs with a 40 mm diameter and 0.1 mm gap. Initially, a time sweep experiment was done at curing temperature (60°C) to check the crosslinking effect of both the film-forming solutions (0.3 mL of Control and PGL-0.7) over time at an angular frequency of1 Hz and 1% strain. The change in complex modulus (G_i_), and complex viscosity (η_i_) was plotted against time (sec). This kinetic study helps to understand the time required for crosslinking (sol-to-gel transformation) at a fixed temperature.

After the kinetic study, the control and PGL-0.7 gels underwent a strain sweep experiment in the range of 10–1000% at 1 Hz to determine the linear viscoelastic region (LVR) and the strain at which the gels were deformed (gel-to-sol transformation).

Frequency sweep test was done at a fixed strain of 10% below the LVR region and in the frequency range of 0.1–10 Hz at 25°C to study the effect of angular frequency (ω) on storage modulus (G’) and loss modulus (G”). The power law model was fitted by using origin software on G’ and G” vs ω plots and a’, a” (pre-exponential factors describing the strength of hydrogels) and b’, b” (power law exponents describing the dependency on angular frequency) were determined (Cui et al., 2022).

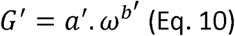

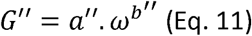

### 2.15. Thermal stability

The thermal stability of the PGL and control films was studied by a Thermogravimetric analyzer (TGA) using the TA SDT Q600 model at 10°C.min^−1^ heating rate between 30-800°C. Limiting Oxygen Index (LOI) of the films was measured using equation 12 to assess their flammability.

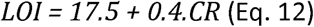

Where CR is the char residue of the films at 800°C (Brenner et al., 2020).

### 2.16. Biodegradation study

Biodegradation study utilizing both hydrolytic and soil burial degradation methods, was done on PGL (PGL-0.7), control, and commercial LDPE films as per Katakrit Chaisuwan et al. (CHAISUWAN et al., 2023).

For the hydrolytic degradation assessment, each film sample was cut into 2 × 2 cm^2^ and immersed in 50 mL of water for 30 days. The initial mass of the films (W_1_) and the final mass of dried film samples after 30 days (W_2_) were measured. Hydrolytic degradation percentage was calculated using Eq. 13.

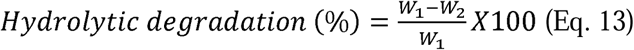

Soil burial test was conducted in lab environment to evaluate the biodegradation of PGL-0.7, control, and commercial LDPE films Soil samples were taken from the Department of Biotechnology, IIT Hyderabad campus (latitude and longitude 17.59499926778957, 78.12314261088649, respectively) and sieved to achieve a consistent particle size of 1 mm.

Initially, 100 g of the prepared soil was added to 7 cm test containers. Subsequently, three film samples, PGL, control, and commercial LDPE each measuring 2 × 2 cm², were placed on the soil surface and covered with an additional 50 g of soil. This experiment was carried out for 90 days, and 3 mL of water was sprinkled on the soil every five days to maintain moisture levels. The films were removed from the soil after 30, 60, and 90 days, and brushed to remove soil from the surface. The initial mass of the films (W_i_) and the final mass of degraded film samples (W_f_) after each 30-day interval were measured. The soil degradation percentage was then calculated by measuring the weight loss using Eqn. 14:

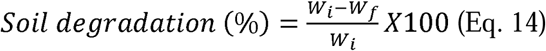

### 2.17. Fruit preservation studies

Fresh Kinnow oranges were bought from a local store of IIT Hyderabad (latitude and longitude 17.584267385597144, 78.11996373588407, respectively). Oranges were packed with control, PGL films (PGL-0.7) and commercial LDPE films and placed at room temperature for 7 days (Fig. 12A). Different physical and chemical properties like weight loss percentage (%), firmness (kg), pH, titrable acidity (TA), malondialdehyde content (nmol/g), total soluble sugar (g), antioxidant capacity (%), and ethylene production by kinnow oranges are tested before and after 7 days.

#### 2.17.1. Weight loss percentage

Kinnow oranges were weighed at the beginning and after 7 days of storage duration to evaluate the weight loss percentage using the Eq.15.

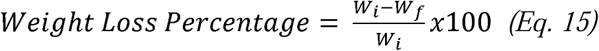

where W_i_ is the initial weight of the fruit, and W_f_ is the fruit weight after 7 days.

#### 2.17.2. Fruit Firmness

The firmness of stored oranges was assessed using a LABART Fruit Hardness Tester, a handheld device fitted with an 8-mm cylindrical probe. Firmness measurements were conducted at five distinct locations of the oranges, with each sample being tested in triplicate (Bodaghi, 2024). The firmness values are calculated in kilograms (Kg). The readings initially recorded in kg/cm^2^ were multiplied by the probe’s surface area (0.5 cm^2^) to obtain fruit firmness values.

#### 2.17.3. Fruit pH and TA

The pH of orange fruit samples was determined using an Eutech CyberScan pH meter. To prepare the samples, 50 g of orange pulp was homogenized and centrifuged at 5000 rpm for 20 minutes. The clear supernatant obtained after centrifugation was used for pH measurement (Bodaghi, 2024).

The Titrable acidity indicates the amount of citric acid in the juice. To fresh juice diluted 10-fold with boiling water (100 mL), 3-4 drops of phenolphthalein indicator were added. The solution was titrated against 0.1 N NaOH until the pink endpoint appeared. The total titrable acidity was measured using Eq. 16:

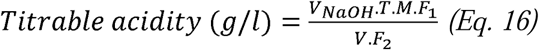

Where, V_NaOH_ =volume of NaOH consumed, T= Concentration of NaOH, M =Molecular weight of Citric acid (192.13 g/mol), V= volume of sample=25 mL, F_1,_ and F_2_ are conversion factors (F_1_=1, F_2_ = 3 as citric acid is a tri-basic acid).

#### 2.17.4. Lipid peroxidation

To determine the malondialdehyde (MDA) content of kinnow orange, the method of X. Li et al. was followed (X. Li et al., 2011). Tissue samples (2 g) were frozen in liquid nitrogen, rapidly ground, and extracted in 5 ml of 10% (w/v) trichloroacetic acid (TCA). The homogenized samples were then centrifuged at 5000 rpm for 20 min. The supernatant (2 mL) was mixed with 2 mL of 10% (w/v) TCA containing 0.5% (w/v) thiobarbituric acid (TBA) solution. The mixture was heated at 100 °C for 20 min, then quickly cooled and centrifuged at 5000 rpm for 10 min. Absorbance was measured at 532 and 600 nm using a Jenway UV/Vis spectrophotometer, and the MDA content was calculated according to the provided Eq. 17.

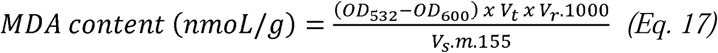

V_t_, V_r,_ and V_s_ were the total volume of the extract and reaction mixture solution, and the volume of the extract solution used in the reaction, respectively, and m was the mass of samples.

#### 2.17.5. Fruit’s Antioxidant Capacity

The antioxidant capacity of the orange extract was assessed using the 2,2-diphenylpicrylhydrazyl (DPPH) radical scavenging assay (Eshghi et al., 2022). This method combined 10 μL of the orange extract with 3 mL of a 0.1 mM DPPH solution. The resulting mixture was thoroughly shaken and incubated in the dark for 1 hour. After the incubation period, the reduction in absorbance was measured at 517 nm using a Jenway UV/Vis spectrophotometer. The antioxidant capacity of the orange extract was subsequently calculated based on the absorbance values obtained from the measurements using Eq. 18. The DPPH assay is widely used to evaluate free radical scavenging ability of various substances, including plant extracts. It provides insight into their potential health benefits due to antioxidant properties.

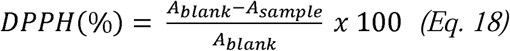

A_blank_ is the absorbance of the control sample, and A_sample_ is the absorbance of the sample.

#### 2.17.6. Sucrose content

To quantify the sucrose content, a standard solution was prepared by combining 1.0 mL of 100-ppm sucrose solution, 1.0 mL of 5% phenol, and 1.0 mL of distilled water, followed by stirring for 1 min. Subsequently, 5.0 mL of concentrated sulfuric acid (H□SO□) was added to the mixture, which was then shaken for 3 min. The solution was allowed to settle for 30 min and cooled in water for 20 min before measuring the absorbance at 490 nm using a UV-Vis spectrophotometer. A blank sample was prepared following the same procedure, omitting the sucrose. A standard calibration curve was plotted using known sucrose concentrations.

From the orange juice sample diluted to 0.050 mg/mL, 1 mL aliquot was combined with 1.0 mL of 5% phenol and 1.0 mL of distilled water. Further, 5.0 mL of concentrated H□SO□ was added, and mixed for 3 min. The solution was allowed to settle for 30 min, cooled in water for another 20 min, and absorbance was measured at 490 nm using a UV-Vis spectrophotometer. A blank sample was also prepared using the same procedure without the orange juice. The sucrose content (%) was determined using the standard calibration plot (Lam et al., 2021).

#### 2.17.7. Ethylene gas production

Ethylene gas production due to the overripening of packaged and unpackaged oranges before and after 7 days was monitored using a handheld ethylene gas sensor (Ambetronics Engineers Pvt. Ltd., Model No.: PG-100-SL). Syringe was attached to the orifice of the probe, and secured with tape. Oranges were kept inside an airtight Ziplock bag, with a PTFE/Silicon septum attached, where a needle could be penetrated several times without puncturing the bag. Ethylene gas concentration (ppm/g) in the headspace was checked every hour after 7 days till 7h.

## 3. Results and discussion

### 3.1. Regression analysis of Lignin extraction from cotton stalk by acid alkali method

The lignin yield from cotton stalk (CS) was significantly affected by the reaction conditions of the acid-alkali process. To enhance the lignin yield and purity, a design of experiments (DOE) based on the Plackett-Burman method was implemented, considering six factors at two levels each.

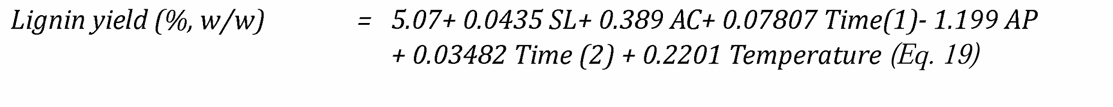

Using the data summarized in Table 1, a regression model (Eq. 19) was developed to predict lignin yield (%), utilizing Minitab 17 software. The regression equation incorporates the independent factors (or variables) A, B, C, D, E, and F which represent the solid loading (SL%, w/v), sulfuric acid concentration (AC %, v/v), acid pretreatment time (time 1, min), alkali concentration (%, w/v), alkali treatment time (time 2, min), and temperature (°C), respectively. While the lignin yield (%, w/w) is the response (dependent) variable.

The first term in Equation 19 represents a constant (5.07), whereas the coefficients associated with each variable correspond to their respective predictor values as determined by the Plackett-Burman (PB) model. Based on Equation 19, it is evident that the solid loading (%, w/v), sulfuric acid concentration (%), acid pretreatment time (min), alkali treatment time (min), and temperature exhibit a directly proportionality with lignin yield (%) whereas alkali concentration (%) demonstrates an inverse relation with lignin yield.

The adjusted R² (76.85%) and predicted R² (70.44%) values supported the model’s reliability, indicating a strong predictive capability. The analysis of variance (ANOVA) for lignin yield is presented in Table 2, which shows the p-value of lack of fit > 0.05, making it insignificant and the model significant.

**Table 2.**
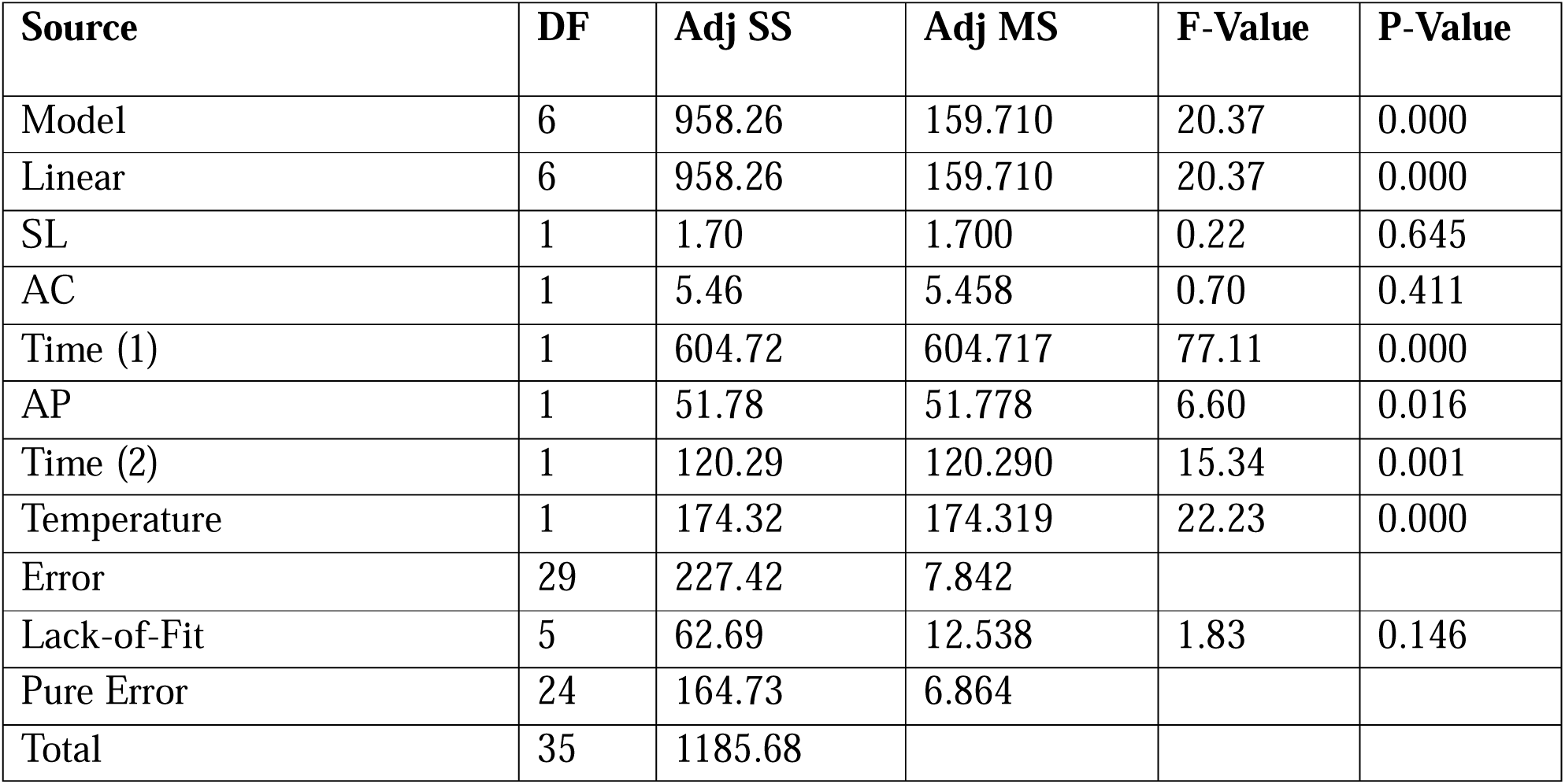
Analysis of Variance using Plackett-Burman for lignin extraction process.

| **Source** | **DF** | **Adj SS** | **Adj MS** | **F-Value** | **P-Value** |
| --- | --- | --- | --- | --- | --- |
| Model | 6 | 958.26 | 159.710 | 20.37 | 0.000 |
| Linear | 6 | 958.26 | 159.710 | 20.37 | 0.000 |
| SL | 1 | 1.70 | 1.700 | 0.22 | 0.645 |
| AC | 1 | 5.46 | 5.458 | 0.70 | 0.411 |
| Time (1) | 1 | 604.72 | 604.717 | 77.11 | 0.000 |
| AP | 1 | 51.78 | 51.778 | 6.60 | 0.016 |
| Time (2) | 1 | 120.29 | 120.290 | 15.34 | 0.001 |
| Temperature | 1 | 174.32 | 174.319 | 22.23 | 0.000 |
| Error | 29 | 227.42 | 7.842 |  |  |
| Lack-of-Fit | 5 | 62.69 | 12.538 | 1.83 | 0.146 |
| Pure Error | 24 | 164.73 | 6.864 |  |  |
| Total | 35 | 1185.68 |  |  |  |

**Table 3.** Mechanical properties of control and PGL films.

| **Sample** | **Ultimate Tensile stress (MPa)** | **Elongation at Break (%)** | **Toughness (MPa)** |
| --- | --- | --- | --- |
| **Control** | 34.62 | 1.94 | 0.293 |
| **PGL-0.5** | 37.7 | 2.2 | 0.329 |
| **PGL-0.7** | 48.5 | 3.1 | 0.905 |
| **PGL-0.9** | 20.91 | 1.45 | 0.102 |
| **PGL-0.7 (300 μL plasticizer)** | 42.67 | 8.1 | 2.554 |
| **PGL-0.7 (350 μL plasticizer)** | 16.47 | 17.4 | 2.444 |
| **PGL-0.7 (400 μL plasticizer)** | 8.55 | 53.1 | 4.708 |
| **PGL-0.7 (450 μL plasticizer)** | 8.5 | 72.1 | 5.681 |
| **PGL-0.7 (500 μL plasticizer)** | 8.4 | 93.9 | 6.515 |

The Half-Normal Plot (Fig. 2A) illustrates the standardized effects of various factors on the response variable, Lignin Yield (%, w/w).

Significant effects are highlighted in the plot with red squares, while non-significant effects are marked with blue circles. Acid pretreatment time (min), alkali concentration (%, w/v), alkali treatment time (min), and temperature (°C) showed significant effect on lignin yield. While the solid loading (%, w/v) and sulfuric acid concentration (% v/v) exhibited insignificant effects.

The Pareto Chart (Fig. 2B) further illustrates the standardized effects of various factors on lignin yield %, highlighting that C (acid pretreatment time), D (alkali concentration), E (alkali treatment time), and F (temperature) have significant influence. Among these, acid pretreatment time was identified as the most critical factor, as increasing its duration enhanced the removal of hemicellulose bound to lignin and cellulose, thereby improving lignin yield. Similarly, higher temperatures and prolonged alkali treatment effectively disrupted hydrogen bonds, facilitating lignin extraction. However, increased alkali concentration may lead to lignin degradation, reducing both its purity and yield. This observation aligns with Equation 19, where alkali concentration exhibited an inverse relationship with lignin yield. The results from the Pareto Chart are consistent with those from the Half-Normal Plot, reinforcing the significant roles of acid pretreatment time, alkali concentration, alkali treatment time, and temperature in maximizing lignin extraction. The chart, arranged in descending order of standardized effects, indicated a statistical significance level (α) of 0.05. Comprehensive understanding of these key parameters is essential for refining biomass processing techniques to enhance lignin recovery for various applications.

Based on these findings, the best conditions for maximum lignin extraction from Cotton stalk were deciphered to be: 5% (w/v) solid loading, 3% (v/v) sulfuric acid concentration, 120 min acid pretreatment time, 3% alkali concentration, 120 min alkali treatment time, and 120°C temperature. Under these conditions, the lignin yield was found to be 48.22 ± 2.45% (w/w).

### 3.2. Pullulan oxidation

Pullulan was oxidized to Pullulan dialdehyde (PDA), which can increase the reaction site by addition of aldehyde groups. The assessment of degree of oxidation by the UV/Vis method quantifies the periodate consumption during oxidation, which was found to be 47.2%. The DO does not necessarily correspond directly to the aldehyde percentage, as hemiacetals and carboxyl groups may also form. Thus, further determination of aldehyde percentage and content (mmol/g) is required, which can be fulfilled by the titration method. The titration is based on the reaction between one equivalent each of hydroxylamine hydrochloride and the aldehyde group of PDA to give one equivalent of hydrochloric acid. This is titrated with NaOH to determine the amount of aldehyde content and percentage. The titration result showed that the aldehyde percentage and content were 20.54% and 1.26 mmol/g, respectively, thus suggesting the presence of hemiacetal groups (Bruneel & Schacht, 1993).

### 3.3. Molecular interactions in PGL films

Pullulan was subjected to oxidation to generate pullulan dialdehyde (PDA) by introducing aldehyde functional groups, which can enhance its reactivity. These aldehyde groups can readily react with the amine groups of gelatin forming stable imine bonds that can contribute to the structural integrity of the biocomposite film (Fig. 2A). Lignin, characterized by its complex polyaromatic architecture, exhibits limited solubility in green solvents due to multiple hydroxyl (−OH) and methoxy (−OCH□) groups. The incomplete solubilization of lignin can result in poor dispersion within biocomposites, primarily due to weak intermolecular interactions. To facilitate effective incorporation, the hydroxyl groups of lignin must be ionized to improve its compatibility with other polymeric components. Beyond electrostatic interactions, hydrogen bonding between the oxygen of lignin’s hydroxyl groups and the hydrogen of gelatin’s amine (−NH□) groups and vice versa can also play a crucial role in stabilizing the lignin-PDA/gelatin matrix.

In this study, lignin was converted into a more reactive form through ionization by dissolving it in a 0.1 N NaOH solution (Oriez et al., 2020). The hydroxyl (−OH) groups of lignin were deprotonated to form oxide (−O□) ions, which were further stabilized by resonance across the polyaromatic backbone. These negatively charged oxide ions established strong electrostatic interactions with the ammonium ions of gelatin (Fig. 2A), thus enhancing the resulting biocomposite film’s overall stability and water resistance.

### 3.4. Chemical analyses and structural insights

#### 3.4.1. 2D HSQC NMR analysis of extracted lignin

2D HSQC NMR is a powerful technique for analyzing lignin’s structure, subunits, and chemical bonds more effectively than ¹H and ¹³C NMR alone (DOI:10.1038/s41598-017-00711-w). The spectrum is divided into three main regions (W. Wang et al., 2020) (A. Kumar et al., 2023) (Fig. 4A(i-iii)) namely, aliphatic side-chain region (δC/δH: 10-40 ppm / 0.2-2.0 ppm), aliphatic oxygenated region (δC/δH: 40-100 ppm / 2-5.6 ppm) (Fig. 4A(i)), and aromatic region (δC/δH: 100-150 ppm / 6-8.5 ppm) (Fig. 4A(iii)).

**Fig. 4.**
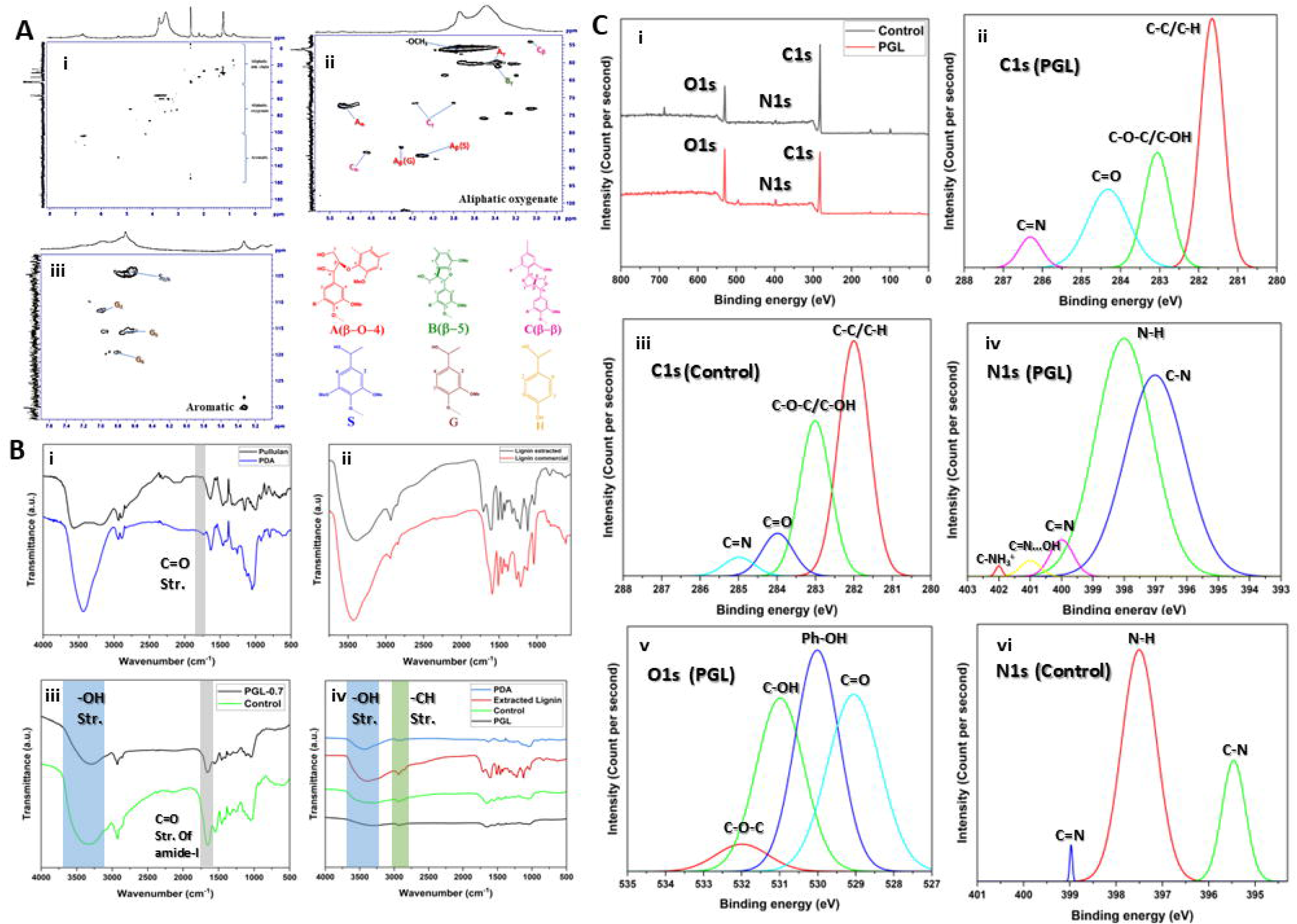
(A)(i-iii) 2D HSQC NMR spectra of extracted alkali lignin; (B) FTIR spectra of (i) pullulan & PDA, (ii) lignin extracted & commercial, (iii) PGL-0.7 & Control films, (iv) PDA, extracted lignin, control, and PGL film; (C) XPS spectrum of PGL (PGL-0.7 with 450 μL 50% glycerol) and control films showing (i) wide spectra (0-800 eV), (ii) C1s peaks, (v) O1s peaks, and (iv) N1s peaks of PGL films and (iii) C1s peaks and (vi) N1s peaks of control films after deconvolution from high-resolution XPS spectra.

The solvent DMSO-d peak is observed at δC/δH 40/2.5 ppm (Fig. 4A(ii)) (A. Kumar et al., 2023).

In the aliphatic oxygenated region, the peak at δC/δH 55.6/3.73 ppm corresponds to the - OMe group, a predominant functional group in lignin. Peaks related to resinol (β-β’) substructures, which are stable under alkaline conditions, appeared at δC/δH 54/3.1 ppm, 63/3.35 ppm, and 71/4.1 & 3.8 ppm for Cβ, Bγ, and Cγ, respectively. The β-O-4’ substructure, another major lignin unit, was observed at δC/δH 71.8/4.8 ppm for Aα. The Cβ-Hβ correlations at δC/δH 83.4/4.27 ppm and 85.7/4.09 ppm correspond to G (Aβ(G)) and S (Aβ(S)) units, respectively (C. Wang et al., 2017). The phenyl coumarin (β-5) substructure was detected at low levels, with Cα signals at δC/δH 86.8/4.6 ppm (C. Wang et al., 2017).

In the aromatic region (Fig. 4A(iii)), peaks for Syringyl (S) and Guaiacyl (G) subunits were present, while the p-hydroxyphenyl (H) subunit was not detected (T. Zhang et al., 2023) (C. Wang et al., 2017). The Guaiacyl (G) signals appear at δC/δH 111/6.9, 115/6.7, 116/6.8, and 119/6.7 ppm for G2, G5, and G6, respectively(T. Zhang et al., 2023). The Syringyl (S) subunit was observed at δC/δH 104.3/6.7 ppm for S2,6.

#### 3.4.2. FTIR analysis

The FTIR spectra of PDA, extracted lignin, control films, and PGL films are illustrated in Fig. 4B. The broad absorption band observed at 3420 cm□¹ in PDA and lignin corresponds to O-H stretching vibrations (Fig. 4B(i,ii)). This band exhibited higher intensity in extracted lignin, likely due to its elevated hydroxyl group content (Ab Rahim et al., 2018). In the control and PGL films, the O-H stretching band shifted to 3328 cm□¹ and 3348 cm□¹ due to its overlapping with the N-H stretching vibrations of amide A. The C-H stretching vibration was detected at 2936 cm□¹ in both PDA and the control film, whereas in lignin and PGL films it appeared at 2925 cm□¹ (Fig. 4B(1-4)). A minor peak at 1726 cm□¹ in PDA corresponds to C=O stretching vibrations (Roy et al., 2023) (Fig. 4B(i)); however, this peak was absent in the control and PGL films, indicating imine bond formation. A distinct peak at 1705 cm□¹ in extracted lignin represents C=O stretching vibrations (Fig. 4B(ii)). The band at 1659 cm□¹ corresponds to the C=O stretching vibration of amide I for both the control and PGL films (Liu et al., 2021) (Fig. 4B(iii)). The C=N stretching frequency of imine bonds overlaps with the amide I C=O stretching, preventing the appearance of a separate C=N peak (L. Zhang et al., 2019). At 1535 cm□¹, amide II-associated C-N bending and stretching vibrations were observed in both the control and PGL films. C-O stretching vibrations were present in all four samples, which attribute to abundant hydroxyl functional groups. Furthermore, at 1237 cm□¹, N-H bending and C-N stretching vibrations characteristic of amide III were evident in the control and PGL films (L. Zhang et al., 2019) (Liu et al., 2021). A progressive decrease in the intensity of the O-H stretching band was noted in PDA and lignin relative to the control and PGL films, which suggested secondary interactions involving hydroxyl groups. Between the control and PGL films, the O-H band was less intense in PGL films, likely due to hydrogen bonding and electrostatic interactions between the hydroxyl groups of lignin and the amine groups of gelatin. These spectral findings confirmed the presence of both chemical and physical interactions among the PGL film components.

#### 3.4.3. XPS analysis

The elemental composition of the PGL and control hydrogel was analyzed, as depicted in Fig. 4C(i). The deconvoluted C1s spectrum of the PGL hydrogel (Fig. 4C(ii)) revealed four distinct peaks, i.e., 281.65 eV corresponding to C-C or C-H bonds, 283.06 eV corresponding to C-O-C or C-OH groups, 284.3 eV corresponding to C=O bonds, and 286.32 eV corresponding to C=N bonds (Afshari et al., 2022). These peaks closely resembled those observed in the deconvoluted C1s spectrum of the control hydrogel (Fig. 4C(iii)). The N1s and O1s spectra were also deconvoluted to further elucidate specific interactions.

In the N1s spectrum, peaks were detected at 398.02 eV for N-H, 397.03 eV for C-N, 399.99 eV for C=N, 401.01 eV for C=N…HO indicating hydrogen bonding, and 402.03 eV for C-NH_3_^+^ suggesting electrostatic interactions (Fig. 4C(iv)) (Stevens et al., 2014) (He et al., 2017). The presence of peaks at 286.32 eV in the C1s spectrum and 399.99 eV in the N1s spectrum confirmed the formation of imine bonds resulting from the reaction between PDA and gelatin (Afshari et al., 2022). Moreover, the peaks at 401.01 eV and 402.03 eV confirmed the existence of both hydrogen bonding and electrostatic interactions between lignin and the gelatin-PDA control hydrogel (Fig. 4C). These interactions were absent in the deconvoluted N1s spectrum of the control hydrogel (Fig. 4C), further validating the interaction between lignin and the PDA/ Gelatin hydrogel.

The deconvoluted O1s spectrum (Fig. 4C(v)) exhibited peaks at 529.03 eV corresponding to C=O bonds, 530.00 eV corresponding to aromatic phenyl hydroxyl groups, 531.01 eV corresponding to C-OH groups, and 532.01 eV corresponding to C-O-C groups. The C=O peaks observed at 284.3 eV in the C1s spectrum and 529.03 eV in the O1s spectrum provided compelling evidence for the conversion of pullulan into PDA, further substantiating the titration results for PDA formation.

### 3.5. PGL films responses to water

PGL films were analyzed for water swelling characteristics and barrier performance in the presence of water. For food packaging materials, controlled water absorption is essential to manage moisture from food or accidental spills without compromising the integrity of the packaged product. Hydrogels exhibit swelling behavior in water due to the presence of both chemical and physical crosslinking. The water molecules can infiltrate the three-dimensional (3D) network structure upon water absorption, leading to expansion similar to other hydrogel-based systems (Fig. 5A). Excessive swelling can lead to film disintegration in wet conditions, adversely affecting mechanical strength and overall usability for food packaging purposes. In the control films, which rely solely on imine-based chemical crosslinking, the swelling ratio reached 143% immediately upon immersion in water and progressively increased to 419% after 1000 mins (Fig. 5B).

**Fig. 5.**
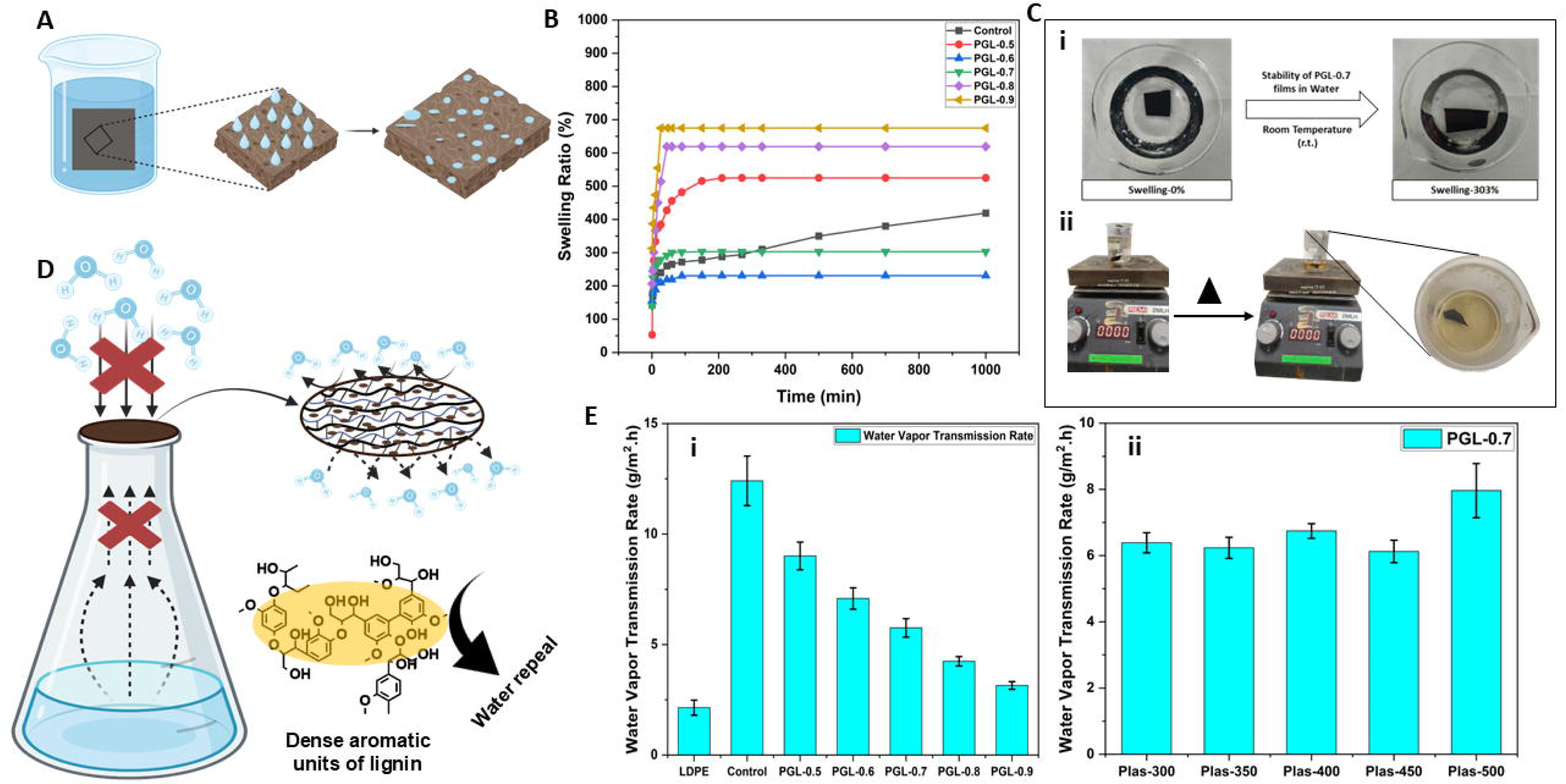
(A) Water swelling behaviour of hydrogel; (B) Water Swelling ratio of control and PGL (0.5-0.9) films over time; (C) Water stability of PGL-0.7 films in both (i) ambient and (ii) hot water; (D) Water vapour barrier mechanism of PGL films; (E) Water Vapour Transmission Rate (g/m^2^.h) at 40±2°C for 24h of (i) control and PGL (0.5-0.9) films and (ii) PGL-0.7 films with varying plasticizer content (300-500 μL 50% glycerol).

Incorporating 0.5 g of lignin into the control films (PGL-0.5) facilitated additional physical crosslinking between the hydroxyl groups of lignin and the amine groups of the film matrix. This reinforcement enhanced the crosslinking density, resulting in more significant swelling than the control films. However, the complex heterostructure of lignin initially impeded water diffusion into the 3D network, leading to a lower initial swelling ratio of 53.24%. Over time, the swelling ratio increased, reaching 525% after 1000 minutes due to the strengthened crosslinking interactions.

Further increasing the lignin content to 0.7 g (PGL-0.7) introduced more hydrophobic interactions within the network, limiting water retention and reducing the swelling ratio to 303%, which was 1.4-fold lower than the control films (Fig. 5C(i)). However, the film’s porosity and brittleness increased at higher lignin concentrations (0.8 g and 0.9 g), due to enhanced hydrogen bonding between lignin and the polymer matrix. This intensified crosslinking, counteracting the hydrophobic effects, that led to increase swelling ratios (PGL-0.8, 619%) and (PGL-0.9, 675%), rendering these films unsuitable for food packaging applications (Fig. 5B). From this result it can be inferred that PGL-0.7 film with a relatively lower swelling ratio can be a potential packaging material even under higher humid conditions.

Water vapor from food and the surrounding environment can significantly impact food shelf life, necessitating food packaging films with a low water vapor transmission rate (WVTR) (Fig. 5D). In comparison, the control films, which consisted of a 3D crosslinked imine network, demonstrated a considerably higher WVTR of 12.41±1.12 g/m^2^.h (Fig. 5E(i)). Although the imine crosslinked structure provided some resistance to water vapor penetration, it is significantly lower than LDPE in limiting moisture transfer.

Incorporating lignin into the control films markedly enhanced their barrier properties by reducing WVTR due to condensed aromatic rings (Lisý et al., 2022). This improvement is attributed to physical and chemical interactions between lignin and the film matrix. The physical interaction occurs primarily through hydrogen bonding between lignin and the PDA/gelatin matrix, while the chemical crosslinks further restrict the free volume available within the film. Together, these interactions impede water vapor diffusion, leading to a reduction in WVTR.

Addition of 0.5 g of lignin to the control films (PGL-0.5) reduced the WVTR to 9.01±0.63 g/m^2^.h. A further increase in lignin content continued to enhance the barrier properties, with PGL-0.7 films exhibiting a WVTR of 5.75±0.42 g/m^2^.h and PGL-0.9 films achieving an even lower WVTR of 3.14±0.17 g/m^2^.h. Among these formulations, the PGL-0.7 films strike the optimal balance between mechanical strength and resistance to water vapor transmission (Fig. 5E(i)).

#### 3.5.1. Effect of plasticizer on WVTR of PGL film

To assess the influence of plasticizers on WVTR, glycerol was incorporated into PGL-0.7 films in varying amounts. The addition of 300 μL of 50% glycerol caused a slight increase in WVTR to 6.38±0.3 g/m^2^.h, while increasing the glycerol content to 450 μL resulted in a negligible change, maintaining a WVTR of 6.13±0.34 g/m^2^.h. However, when the glycerol concentration was increased to 500 μL, the WVTR rose significantly to 7.96±0.82 g/m^2^.h, representing a 1.4-fold increase compared to PGL-0.7 film (Fig. 5E(ii)). The addition of glycerol improved film flexibility; however, excessive amounts compromise moisture resistance, highlighting the need for precise formulation to achieve maximum barrier properties.

### 3.6. Mechanical strength and flexibility of PGL films

Food packaging films must possess high tensile strength and adequate stretchability (or malleability) to ensure durability and resistance to tearing. The control films, composed solely of gelatin and PDA, exhibited a tensile strength of 34.62 MPa but a low elongation at a break of 1.94%, indicating insufficient flexibility for practical use in packaging applications. The incorporation of lignin into the PDA/ Gelatin blend films led to improvements in both tensile strength and elongation at break. Lignin’s heterogeneous structure can enhance the mechanical integrity of the films. However, excessive lignin incorporation can result in aggregation within the matrix, leading to a decline in tensile strength and elongation (Fig. 2B(v-vi)).

Adding 0.5 g of lignin (PGL-0.5) increased the tensile strength to 37.7 MPa and slightly improved the elongation at break to 2.2%. The highest mechanical performance was observed at 0.7 g of lignin (PGL-0.7), where the tensile strength reached 48.5 MPa and the elongation at break improved to 3.1%. However, despite the enhanced tensile strength, the films remained insufficiently malleable for food packaging applications (Fig. 6A). Upon further increasing the concentration of lignin to 0.9g (PGL-0.9) the tensile strength decreased to 20.91 MPa and the elongation at break decreased to 1.45%.

**Fig. 6.**
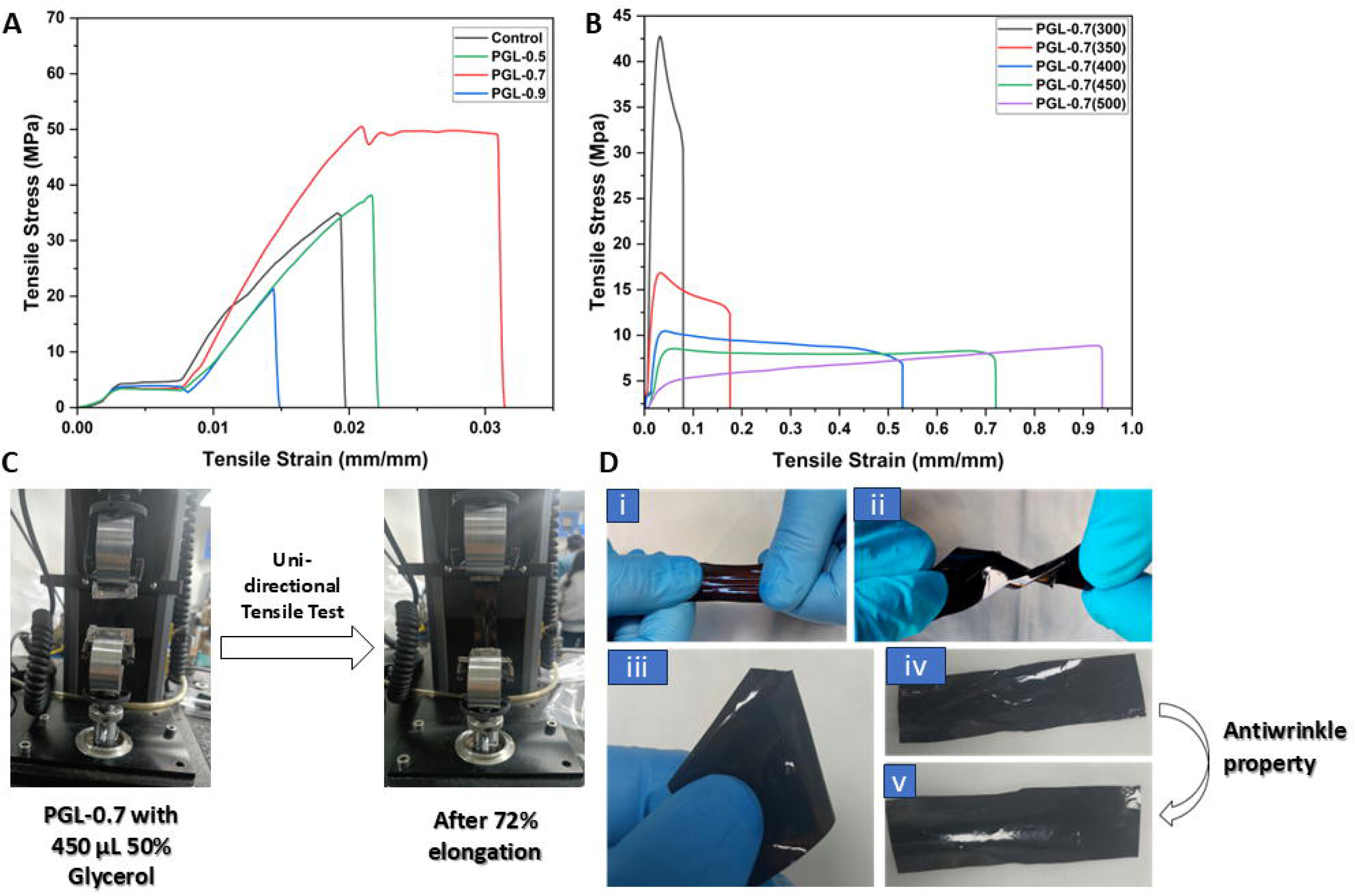
(A) Tensile stress (MPa) vs Strain (mm/mm) curves of control and PGL (0.5-0.9) films, and (B) PGL-0.7 films with varying plasticizer content (300-500 μL 50% glycerol);(C) Uniaxial Tensile testing of PGL-0.7 films with 450 μL 50% glycerol, showing 72% elongation; (D)(i) Stretching of PGL-0.7 films; (ii) Twisting and (iii) Folding of the same film; (iv,v) keeping the same film for some time to show the antiwrinkle effect of the PGL-0.7 film.

#### 3.6.1. Effect of plasticizer on mechanical properties of PGL films

Glycerol was introduced as an external plasticizer, which enhances flexibility and reduces brittleness by interfering with the hydrogen bonds within the polymer matrix by interposing itself between polymer chains, allowing for greater chain mobility and reduced rigidity. Adding 300 μL of 50% glycerol decreased tensile strength to 42.67 MPa while significantly increasing the elongation at break to 8.1%. Despite this modification, the films did not meet the criteria for practical food packaging applications (Fig. 6B). Further increasing the glycerol content reduced tensile strength while progressively enhancing elongation. For instance, incorporating 350 μL of 50% glycerol reduced the tensile strength to 16.47 MPa while increasing the elongation at break to 17.4%. At a glycerol concentration of 500 μL, the tensile strength decreased sharply to 8.4 MPa, whereas the elongation at break increased substantially to 93.9%. Although glycerol can effectively enhance films flexibility, excessive amounts severely compromised mechanical strength, rendering the films unsuitable for food packaging. This study identified 450 μL of 50% glycerol as the ideal concentration to achieve a balanced mechanical performance with a tensile strength of 8.5 MPa and an elongation at a break of 72.1% (Fig. 6C). The final PGL-0.7 film (50% 450 μL glycerol) exhibited exceptional malleability, allowing it to conform easily to various shapes and surfaces (Fig. 6D(i-v)). Additionally, it demonstrated a high elongation capacity, as illustrated in Fig. 6C.

### 3.7. UV blocking properties of PGL films

Ultraviolet (UV) radiation is categorized into three types based on wavelength: UV-A (320– 400 nm), UV-B (280–320 nm), and UV-C (200–280 nm). These rays pose significant risks to human health, with UV-B being the most harmful. UV-B radiation can penetrate food packaging films, leading to food degradation. Additionally, prolonged exposure to UV radiation can accelerate the degradation of synthetic plastics by inducing free radical formation, which cleaves polymer chains and compromises hydrogel integrity. Gelatin, employed as a crosslinking agent in both the control and PGL films, is a protein known for its inherent UV-blocking capabilities, attributed to the presence of aromatic amino acids such as tryptophan and tyrosine (Ezati et al., 2023). While gelatin can effectively block UV-C radiation, its ability to shield against UV-B is limited. Consequently, the control films primarily prevented UV-C penetration but did not provide sufficient protection against UV-B, leaving food items susceptible to photodegradation.

Incorporating 0.5 g of lignin into the control films (PGL-0.5) significantly enhanced their UV-blocking performance. The presence of lignin enables the films to obstruct UV-B radiation completely, along with UV-A and UV-C rays (Fig. 7B&D). This improved UV-shielding ability arises from the chromophoric structures within lignin, which can interact with light to generate quinone derivatives, thereby enhancing UV absorption. Also, an electronic transition within the lignin structure can enhance UV absorption capabilities by binding free electron pairs to aromatic rings. Additionally, π-π stacking of benzene rings within the lignin structure can increase the conjugation by decreasing the HOMO-LUMO gap and enhancing electron transitions to increase UV absorption properties (X. Wu et al., 2024) (Fig. 7A). The inclusion of lignin can reinforce the protective function of the films, effectively preventing food deterioration and extending the durability of the packaging material.

**Fig. 7.**
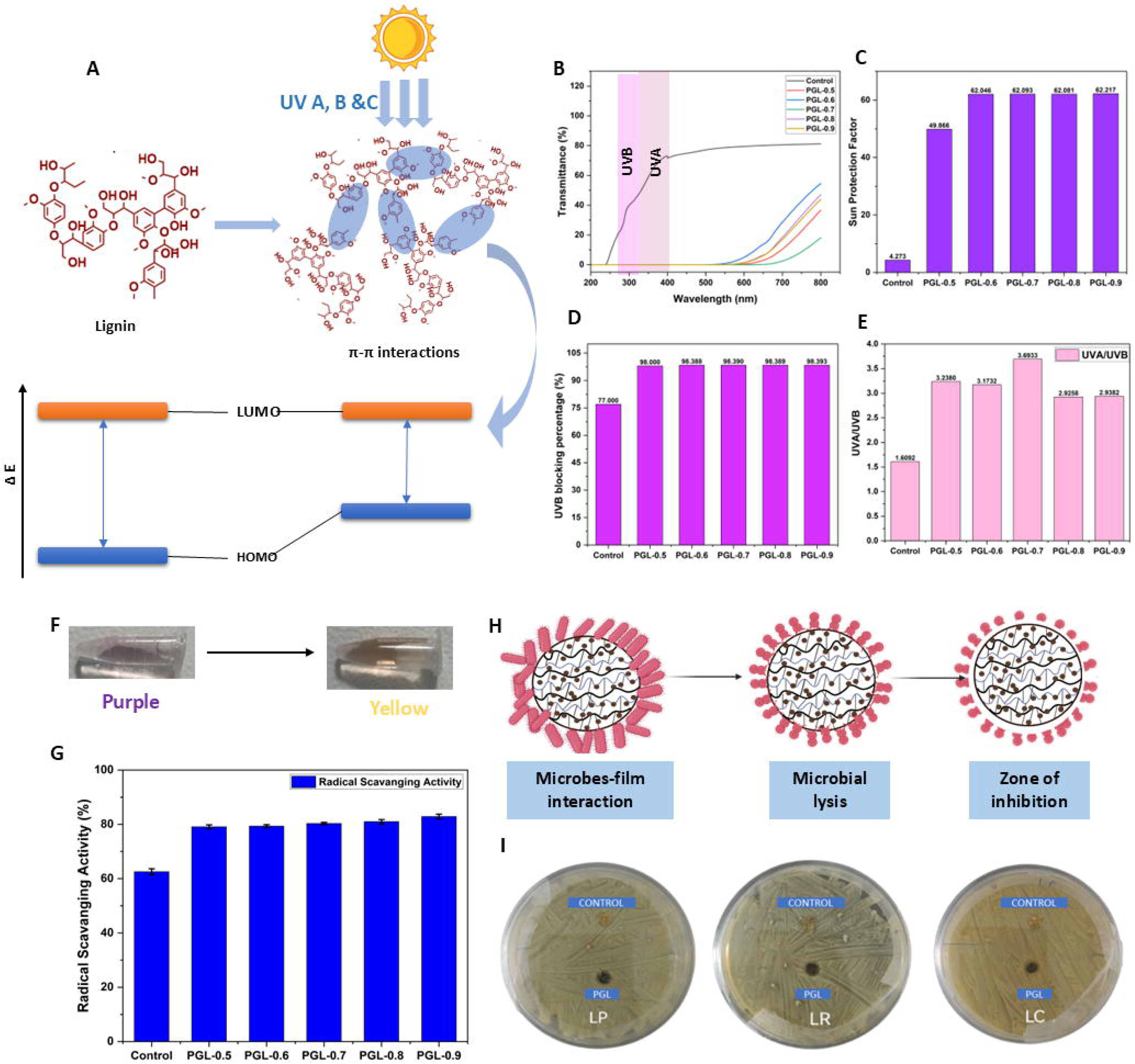
(A) UV absorption mechanism of lignin; (B) UV Transmittance of control and PGL (0.5-0.9) films recorded at 200-800nm; (C) Sun Protection Factor (SPF) and (D) UV-B Blocking Percentage (%) of control and PGL (0.5-0.9) films; (E) UVA/UVB ratio to estimate effectiveness of control and PGL (0.5-0.9) film’s UV absorption properties over broad spectrum; (G) Colorimetric change of DPPH from purple to yellow due to scavenging; (H) Radical Scavenging Activity (%) of control and PGL (0.5-0.9) films; (I) Schematic representation of interaction with antibacterial PGL films with microbes; (J) Antimicrobial activity of control and PGL films (PGL-0.7 with 450 μL 50% glycerol) against *Lactobacillus pentosus* (LP), *Lactobacillus rhamnoses* (LR) and *Lactobacillus casei* (LC) strains.

The corresponding data in Fig. 7C&D presents the Sun Protection Factor (SPF) and UV-blocking efficiency of the control and PGL films. The results indicate that adding 0.5 g of lignin substantially increased the SPF value from 4.2 to 49.86 and enhanced the UV-blocking efficiency from 77% to 98%. A further increase in lignin content to 0.6 g elevated the SPF value to 62.05 and the UV-blocking efficiency to 98.39%. However, additional lignin did not yield significant improvements in SPF or UV-blocking efficiency beyond this concentration. Among the formulations examined, PGL-0.7 demonstrated the most ideal performance, with an SPF value of 62.09 and a UV-blocking efficiency of 98.39%. Additionally, PGL-0.7 films showed effective UV-blocking properties in a broad spectrum, which can be demonstrated by UVA/UVB ratio of 3.69, which is 1.2 folds higher than PGL 0.6 and 2.3 folds higher than control (Fig. 7E). These findings confirmed that incorporating lignin can eliminate the need for synthetic UV filters (Khare et al., 2025), which are typically used in plastic-based food packaging to mitigate UV-induced degradation.

### 3.8. Antioxidant properties

Food components can undergo oxidative reactions, forming free radicals, accelerating food degradation, and significantly reducing shelf life. To mitigate this issue, antioxidant compounds are often incorporated into food products to neutralize these radicals. However, the direct addition of synthetic antioxidants raises potential health concerns due to their possible toxicity. A safer and more sustainable alternative is the development of active food packaging films containing natural antioxidant agents. These films can effectively suppress radical formation, protect food from oxidation, and contribute to an extended shelf life. In this study, gelatin, utilized in the control films, can exhibit inherent antioxidant properties due to its high amino acid content. As a result, the control films demonstrated a radical scavenging activity of 62.54±1.09%. The incorporation of lignin into the film matrix further enhanced this antioxidant activity. Lignin can capture free radicals and further stabilize them by stable delocalization throughout its aromatic structure (Qin et al., 2020) (Fig. S1). Specifically, adding 0.5 g of lignin (PGL-0.5) increased the radical scavenging efficiency to 79.08±0.78%. However, beyond this concentration, the improvement in antioxidant performance became less pronounced. Films containing 0.7 g of lignin (PGL-0.7) exhibited a scavenging activity of 80.3±0.43%, while those with 0.9 g of lignin (PGL-0.9) achieved a marginally higher activity of 82.86±0.89% (Fig. 7G).

The incorporation of lignin significantly enhanced the antioxidant properties of the gelatin/PDA-based control films, with substantial improvements observed at lower concentrations. While increasing lignin content further enhanced radical scavenging activity, the improvement is less significant beyond a certain threshold. This suggests that even minimal lignin addition is sufficient to improve the antioxidant performance of the packaging films. Moreover, this approach provides a sustainable strategy for food preservation by reducing the reliance on synthetic antioxidants. Among the tested formulations, PGL-0.7 was identified as the most effective film, offering comparable antioxidant performance to that of higher lignin-loaded variants (PGL-0.8 and PGL-0.9) while maintaining a balanced composition. Considering low WVTR and high mechanical strength of PGL-0.7 film with glycerol (50% 450 μL glycerol), all the subsequent studies, including the PGL film characterization and application onto kinnow oranges, were conducted using this PGL formulation.

### 3.9. Antimicrobial properties

The antimicrobial efficacy of both the control and PGL films was evaluated against *Lactobacillus pentosus*, *Lactobacillus rhamnosus*, and *Lactobacillus casei*, bacterial strains (Fig. 7H) commonly associated with the spoilage of food items, including fruits. The control film exhibited limited antimicrobial activity, as evident by small inhibition zones. This mild inhibitory effect can be attributed to the presence of amino acids in gelatin, which possess weak antimicrobial properties. The inhibition zones measured for the control film were 0.62 ± 0.28 cm for *L. pentosus*, 0.626 ± 0.03 cm for *L. rhamnosus*, and 0.628 ± 0.14 cm for L. casei. In contrast, the PGL films demonstrated enhanced antimicrobial activity, primarily due to the bioactive properties of lignin. The inhibition zones for the PGL films were recorded as 0.7 ± 0.34 cm for *L. pentosus*, 0.65 ± 0.17 cm for *L. rhamnosus*, and 0.75 ± 0.27 cm for L. casei (Fig. 7I). This result highlights the antimicrobial potential of lignin and its efficacy in enhancing the inhibitory performance of the films against spoilage-associated bacterial strains.

### 3.10. Surface morphology and wettability studies

The surface morphology of the films was examined to inspect any agglomeration or roughness on the films. The surfaces of both the PGL-0.7 and control films were found to be smooth at different resolutions and magnifications, which can be observed from the SEM images (Fig. 8A-D). The surface morphology of the PGL film exhibited no visible agglomeration of lignin or other copolymers, which indicated good miscibility of the polymers within the matrix. The mapping analysis by EDX of the control and PGL films (Fig. 8G&H) prove that there were no toxic impurities that can affect the food packaging applications. A distinct sodium peak was observed for the PGL-0.7 films, as the solvent system used for dissolving lignin was 0.1 N NaOH.

**Fig. 8.**
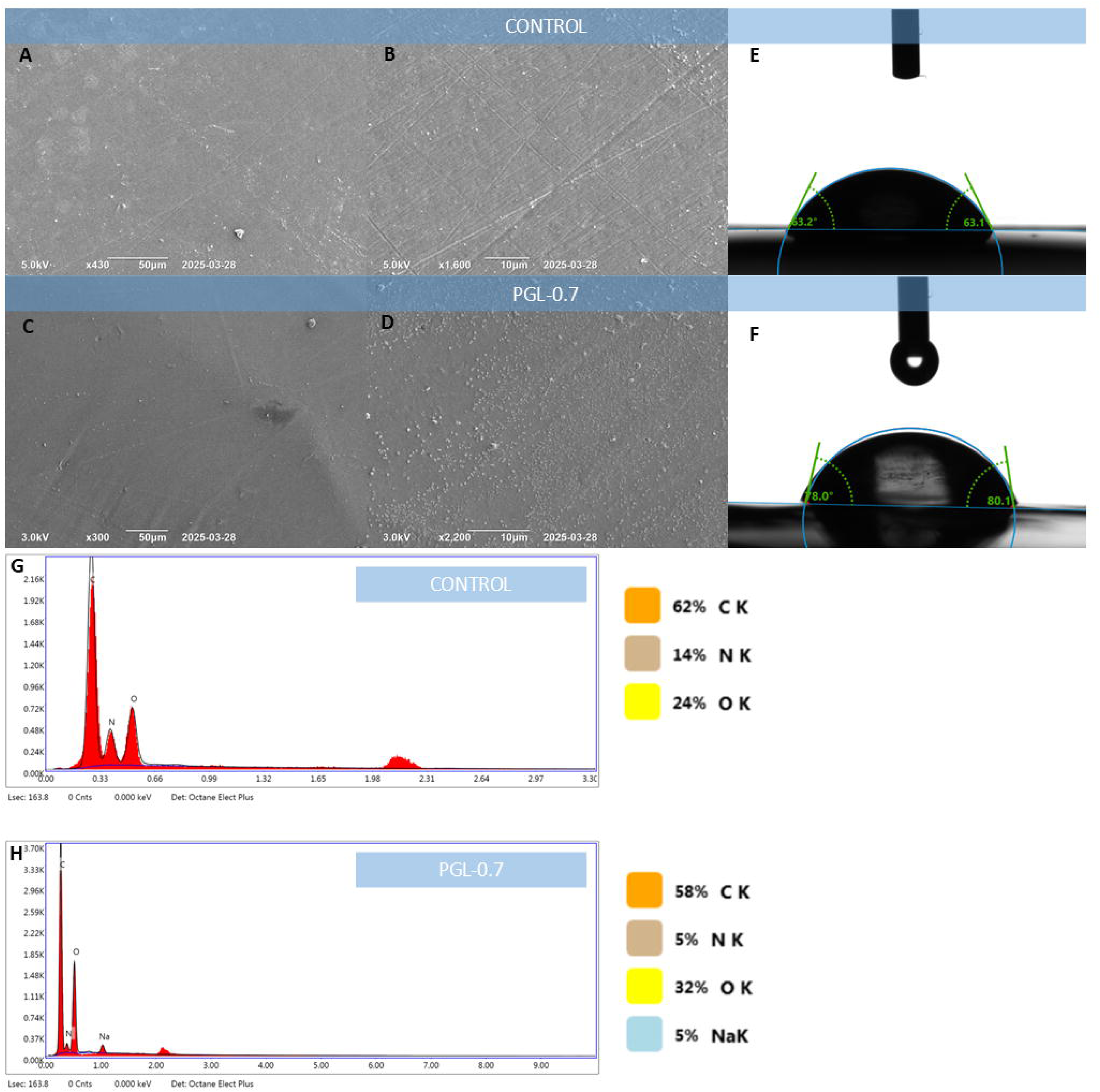
(A &B) SEM images of control & (C&D) PGL-0.7 films; (E) Contact angle measurement of control & (F) PGL-0.7 films; Elemental analysis by EDX of (G) control & (H) PGL-0.7 films.

Surface wetiibility is a crucial factor, which signifies the moisture and microbial resistance of films, essential for effective food packaging. The control film exhibited a contact angle of 63.1 ± 0.050°, while the PGL film demonstrated an increased contact angle of 79.2 ± 0.040 (Fig. 8E&F). The relatively high contact angle in both films can be linked to the smooth surface, as confirmed by SEM images. The contact angle of the PGL film was 1.25-fold higher than that of the control films due to the incorporation of lignin and formation of a denser network structure. From the water swelling ratio, WVTR, and contact angle correlation, it is evident that PGL films exhibit higher values compared to control owing to a smoother surface and higher crosslinked structure.

### 3.11. Hydrogel Network Dynamics and Viscoelastic Properties

Rheological studies were conducted to evaluate the network strength of crosslinked hydrogels and their viscoelastic strength. From the time sweep data, it was inferred that at curing temperature (60°C, isothermal condition), the PGL film-forming solution’s complex modulus and viscosity increased sharply over time. In contrast, in the case of the control film-forming solution, there was no increase in both, highlighting the intense crosslinking in PGL composition at 60°C over time (Fig. 9A&B). An amplitude or strain sweep was done to study the network strength and determine the Linear Viscoelastic Region (LVR), for both the hydrogels G’/G”>1 in the LVR region (Fig. 9C-F), which showed ideal elastic behaviors. The plots revealed that the gel-to-sol transformation occurred at 319% for PGL hydrogel and 119% for control hydrogel, describing the broadness of the LVR region of PGL hydrogel and its high network strength (Fig. 9C&D). Further, a Frequency sweep was conducted, followed by a power law model fitting on the plots in the LVR range to study the hydrogels’ viscoelastic properties (Fig. 9G-J). From the frequency sweep plots, it was observed that the Storage modulus is always greater than the loss modulus in the case of all hydrogels in the LVR region. A high R^2^ value suggests a high degree of power law model fitting on frequency sweep plots (Fig. 9I&J). The low b’ and b” (frequency-dependent factor) values of PGL hydrogel were very low compared to the b’ and b” values of control hydrogels, which described the minor influence of frequency on the inner crosslinked structure of PGL hydrogel (stable networks). The b’/b” value was greater than zero, indicating the presence of both chemical and physical interactions (Cui et al., 2022). The strength of the hydrogels can also be inferred from the pre-exponential values of Eq. 11 (a’). PGL hydrogels showed higher a’ than the controls, substantiating the inner strength of PGL films (Cui et al., 2022).

**Fig. 9.**
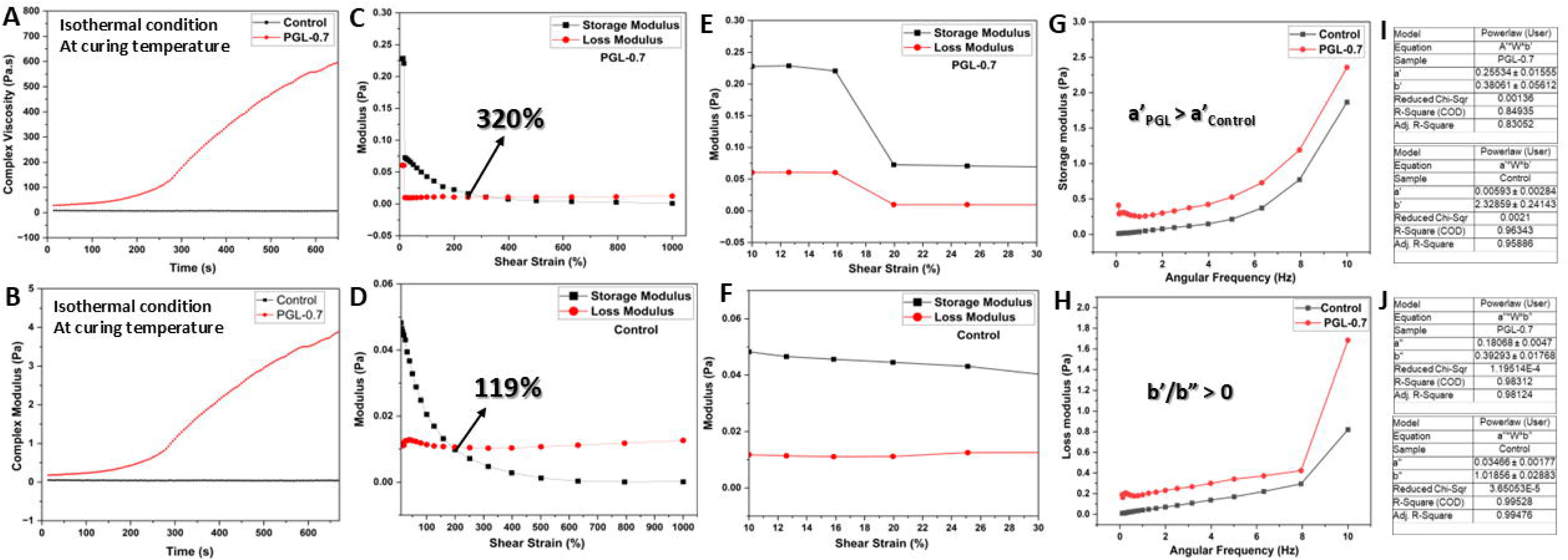
(A&B) Time sweep, (C-F) Amplitude sweep, and (G&H) Frequency sweep with (I&J) Power law model fitting on frequency sweep plots of PGL-0.7 and control film forming solution.

### 3.12. Thermal stability of PGL films

The TGA plot (Fig. 10A) showed that the initial weight loss was due to moisture loss between 50-110°C in the films. In the second stage, between 140-190°C, there was weight loss in both films due to the disruption of peptide bonds of gelatin. The third significant degradation occured at 25.64 and 27.9 % in the case of PGL-0.7 at 270 and 266 °C for control films, respectively which was due to C-C bond decomposition (Fig. 10B). At T> 550 °C, PGL-0.7 films were more thermally stable than the control films, as Lignin can form thermally stable compounds under heat. The LOI of the control and PGL-0.7 films were 26.8 and 27.93, respectively, confirming the thermal stability and non-flammability of the PGL film, as lignin forms an extra char layer (Goliszek et al., 2024), which prevents further ignition (Fig. 10C).

**Fig. 10.**
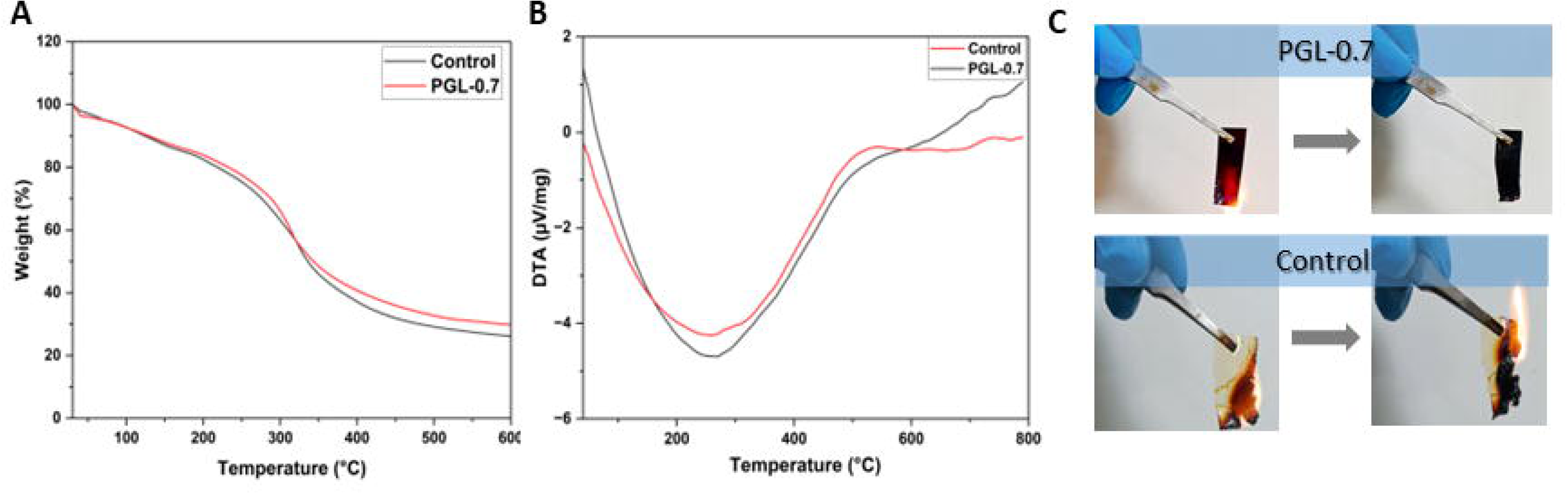
(A) TGA, (B) DTG plots & (C) non-flammability of PGL-0.7 and control films.

### 3.13. Biodegradation ability

The biodegradation behavior of PGL and control films was assessed using hydrolytic and soil burial methods, while commercial LDPE films were included as a reference. The results indicated that the PGL and control films underwent degradation, whereas the LDPE film exhibited no observable degradation, reaffirming its non-biodegradable nature. This resistance to degradation is highly undesirable for packaging applications, as it contributes directly to plastic pollution. Under hydrolytic conditions (Fig. 11B), the control film exhibited a degradation rate of 77.21%, whereas the PGL film degraded to a lesser extent, at 48.37%. In contrast, during the soil burial test (Fig. 11A&C) over 90 days, the PGL film demonstrated a higher degradation rate (40.42%) compared to the control film (35.36%) (Table 4). The commercial LDPE film remained entirely intact in the soil, confirming its persistence in the environment.

**Fig. 11.**
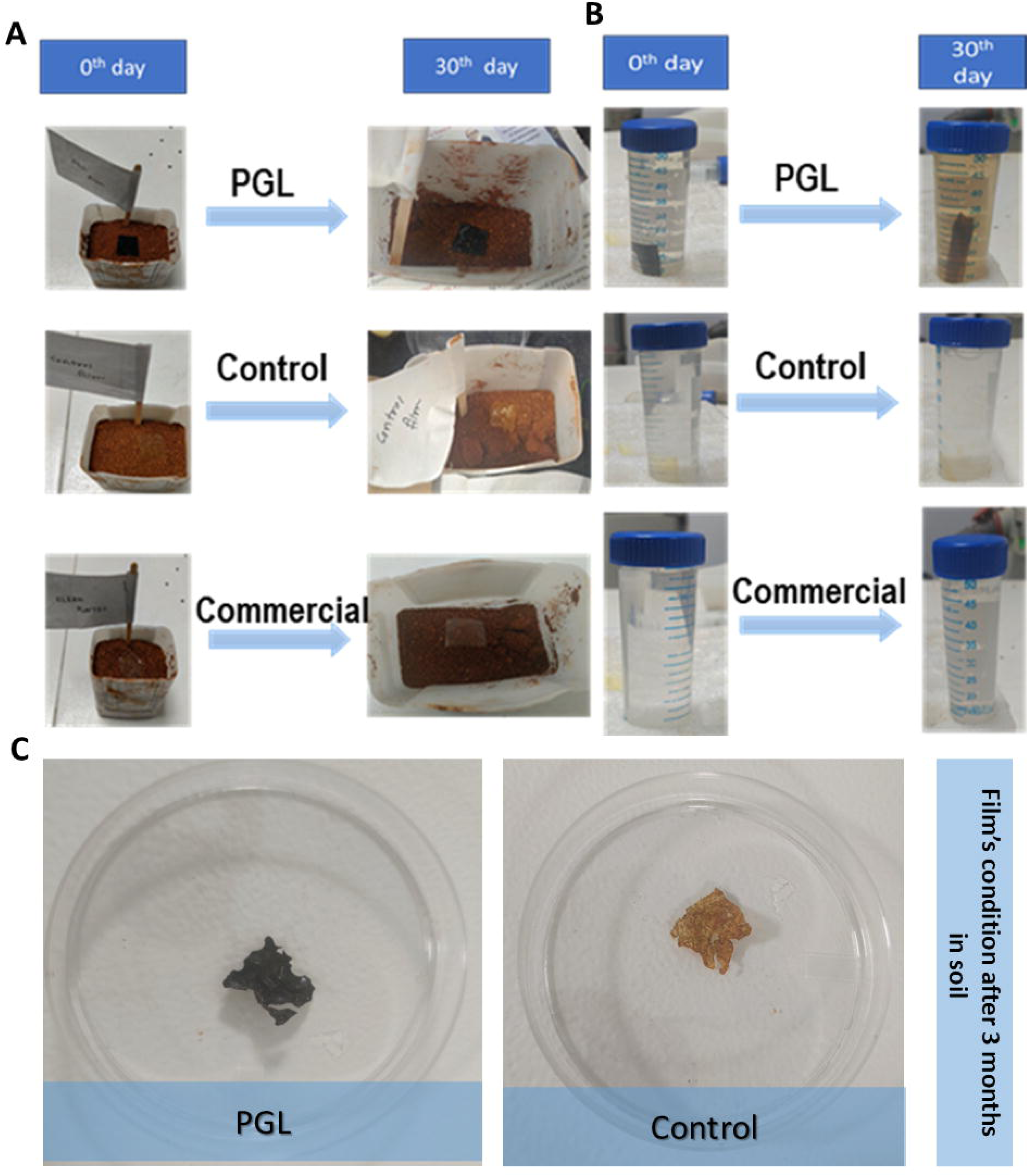
(A&C) Soil Biodegradation of PGL, Control, and Commercial LDPE films over 90 days; (B) Hydrolytic degradation of PGL, control, and commercial LDPE films over 30 days.

**Table 4.** Comparison of biodegradation of PGL, Control, and Commercial LDPE films.

| **Films** | **Hydrolytic degradation (%) after 30 days** | **Soil Degradation (%)** |  |  |
| --- | --- | --- | --- | --- |
|  |  | **30 days** | **60 days** | **90 days** |
| PGL | 48.37 % | 24.71% | 29.7% | 40.42% |
| Control | 77.21% | 12.93% | 23.25% | 35.36% |
| Commercial LDPE films | 0% | 0% | 0% | 0% |

The PGL film exhibited a relatively higher degradation rate under soil burial conditions compared to both the control and commercial counterparts. This suggests that landfill disposal can serve as a viable and convenient method for promoting the biodegradation of PGL films, particularly considering their widespread use in terrestrial environments. This degradation study is currently being continued to investigate for the time required for maximum degradation.

### 3.14. Kinnow orange preservation studies

#### 3.14.1. Weight loss percentage

The weight loss of oranges packaged with PGL, control, and commercial LDPE films, as well as without packaging oranges, was evaluated over a period of seven days (Fig. 12A, B&D). Oranges wrapped in PGL film exhibited a weight loss of 10.75 ± 0.85% (w/w), while those enclosed in the control film showed a weight loss of 10.85 ± 1.08% (w/w). In contrast, oranges packaged with commercial LDPE film experienced significantly lower weight loss, of 0.59 ± 0.089% (w/w). The highest weight loss of 14.52 ± 1.8% (w/w) was observed in unwrapped oranges, as the absence of a protective barrier allowed unrestricted moisture loss. The moisture-absorbing nature of both PGL and control films contributes to the observed weight reduction, as these films absorb moisture released from the fruit. In contrast, due to their dense structure and inherently low WVTR, LDPE films prevent water vapor escape however, lack moisture absorption capability. This characteristic minimizes weight loss in LDPE-packaged oranges and leads to moisture accumulation inside the packaging due to fruit respiration, thereby fostering microbial contamination. Conversely, the moisture-absorbing properties of PGL films help mitigate excessive moisture buildup, thereby improving fruit preservation and maintaining freshness more effectively. Thus, it can be inferred that though the weight loss of PGL packaged fruits is on the higher side compared to LDPE packed fruits, the excellent moisture-absorbing property of PGL films without disintegrating helps in retaining the freshness of the fruit (Table 5 &Fig. 12F).

**Fig. 12.**
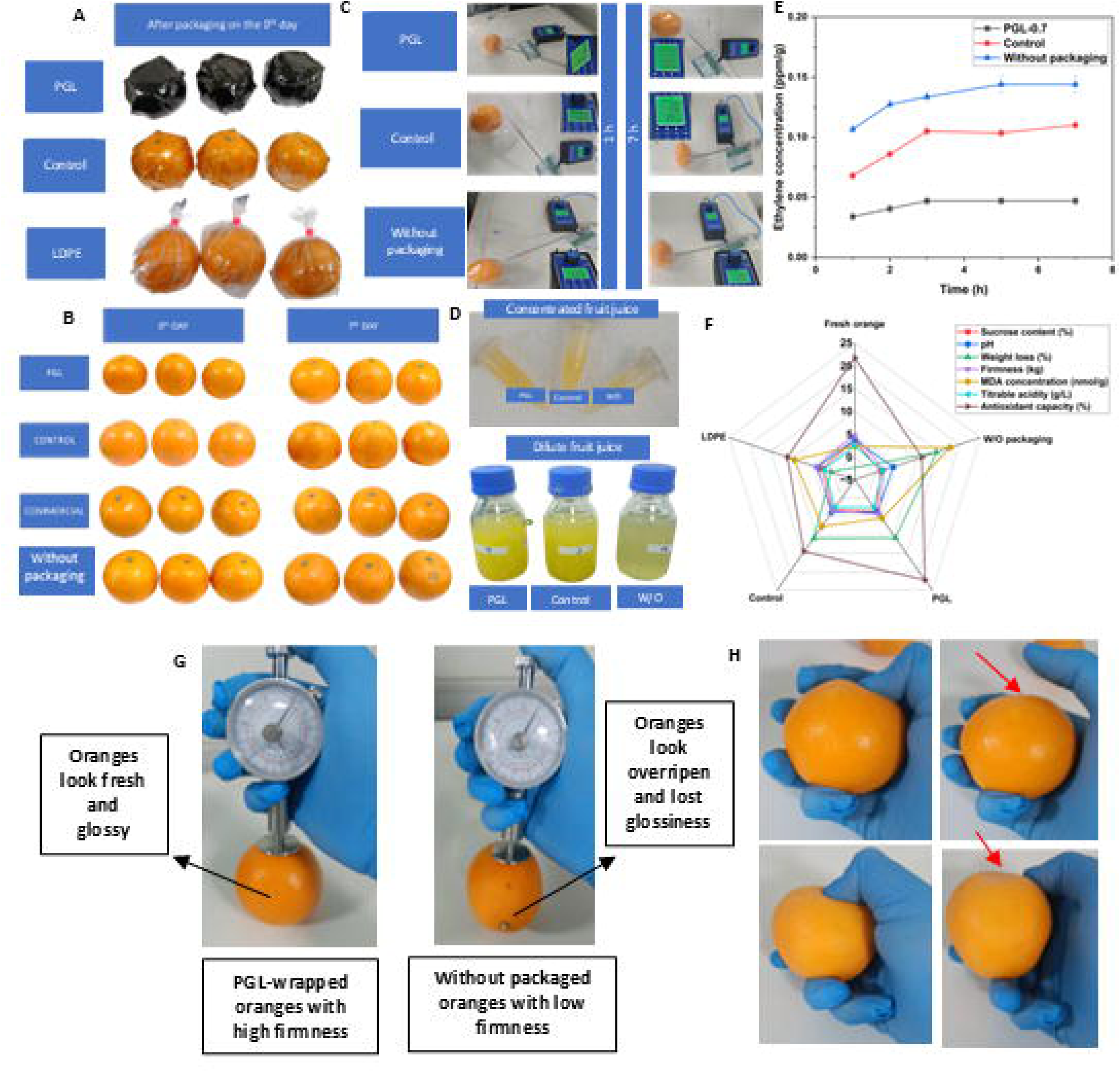
(A&B) PGL, control and commercial LDPE wrapped oranges before and after 7 days; (D) PGL, control and commercial LDPE wrapped orange juice; (C&E) Ethylene gas determination of packed and oranges without packaging by a sensor after 7 days; (F) Spider web plot representing quality assessment of packed and kinnow oranges without packaging; (G&H) Firmness measurement of PGL packed and oranges without packaging.

**Table 5.**
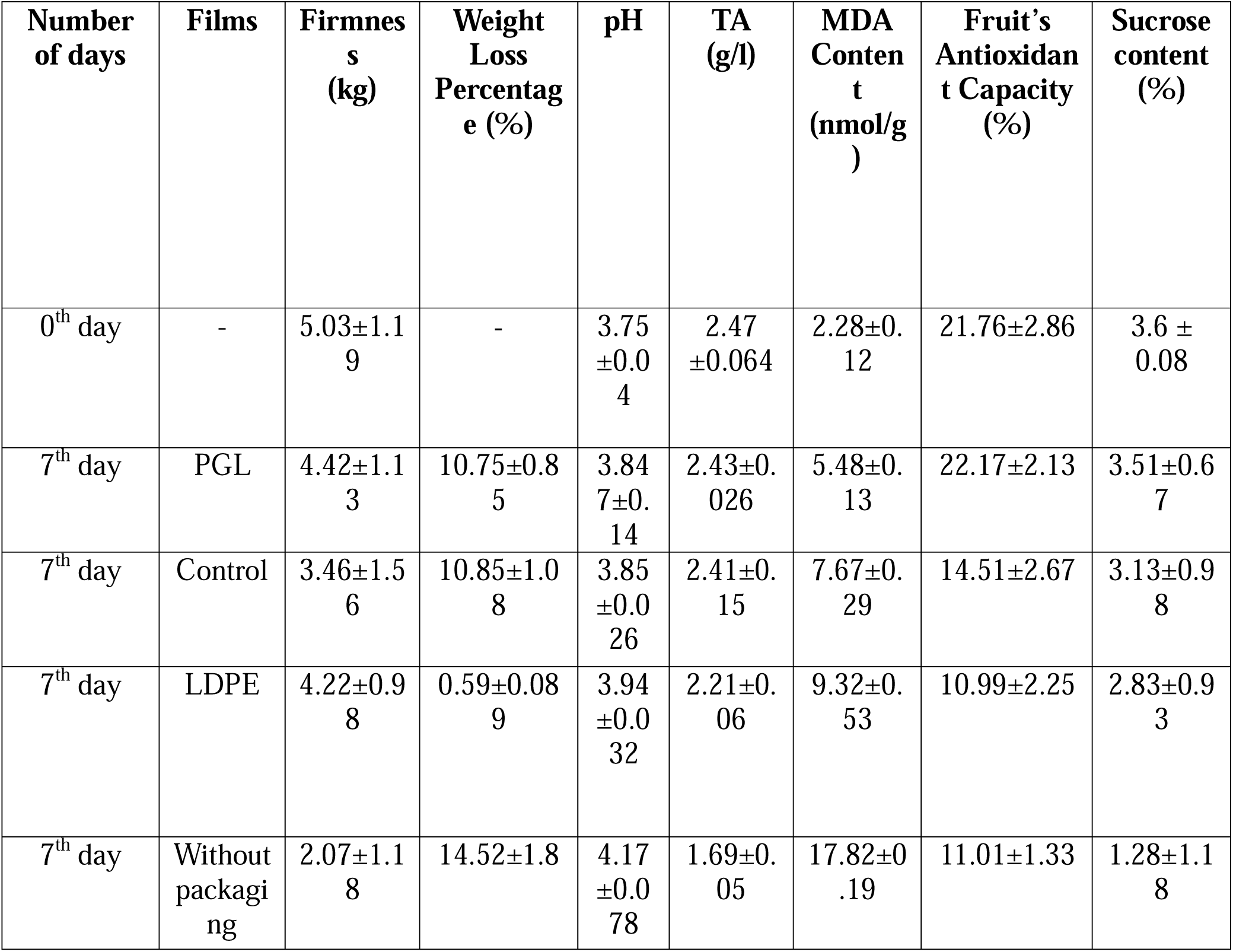
Properties of kinnow oranges before and after 7 days of incubation.

#### 3.14.2. Fruit Firmness

Over time, as moisture is lost and the ripening process advances, the firmness of oranges can decline progressively. On 0^th^ day, fresh oranges had a firmness of 5.03±1.19 kg. In the case of oranges without packaging after 7 days, firmness decreased substantially from 4.74 ± 0.89 kg to 2.07 ± 1.18 kg due to excessive moisture loss and overripening. Oranges wrapped in PGL film, despite experiencing some weight loss, retained a firmness of 4.42 ± 1.13 kg, suggesting that the presence of lignin, an active component in the PGL film, played a role in delaying the ripening process and preserving fruit firmness (Fig. 12G&H).

In contrast, oranges packaged in control film exhibited a more pronounced decline in firmness to 3.46 ± 1.56 kg, coupled with higher weight loss. This indicates the inefficacy of the control films in preventing both moisture loss and overripening, making them less suitable for fruit packaging applications. Meanwhile oranges wrapped in commercial LDPE film experienced minimal moisture loss, and their firmness declined to 4.22 ± 0.98 kg, suggesting that LDPE films are less effective than PGL films in delaying the ripening process (Table 5& Fig. 12F).

#### 3.14.3. pH and TA

During fruit ripening, the pH of the fruit can increase due to the natural degradation of organic acids, such as citric acid, which is a key marker of ripening. TA is inversely related to pH; thus, with the increase in ripening, pH increases and TA decreases. Initially, fresh oranges on day 0 had a pH of 3.75 ± 0.04 and TA of 2.47 ±0.06 g/L. After 7 days for oranges without packaging conditions, the pH rose to 4.17 ± 0.08, and TA decreased to 1.69±0.05 g/L, indicating significant ripening. In contrast, oranges packaged in PGL film exhibited only a slight increase in pH to 3.85 ± 0.14 and a slight decrease in TA to 2.43±0.03, suggesting minimal ripening. Similarly, oranges wrapped in the control film showed pH increase to 3.85 ± 0.03 and TA decrease to 2.41±0.15 g/L. However, oranges enclosed in commercial LDPE film experienced a greater pH rise to 3.94 ± 0.032, indicating a reduction in citric acid and decrease in TA content to 2.21±0.06 g/L. This change was not observed in oranges packaged with PGL or control films, suggesting that LDPE films are less effective in slowing the ripening process than PGL film (Table 5 & Fig. 12F).

#### 3.14.4. MDA concentration

Malondialdehyde (MDA) is a byproduct of lipid peroxidation and can serve as a key indicator of oxidative stress in fruit, ultimately influencing its spoilage which can result from overripening during storage and transportation or exposure to environmental stressors (Stabrauskiene et al., 2025). Lower MDA levels can be correlated with better fruit quality and extended shelf life. Lipid peroxidation in oranges is triggered by producing reactive oxygen species (ROS), which function as free radicals, oxidizing lipids to MDA and accelerating the ripening process (Laguerre et al., 2007) (Fig. S2). To mitigate the effects of these free radicals, the natural antioxidant compounds present in the fruit are gradually depleted. Additionally, prolonged storage can lead to microbial contamination, further exacerbating lipid oxidation (Stabrauskiene et al., 2025).

In the present study, oranges wrapped in PGL film exhibited the lowest MDA levels (5.48 ± 0.13 nmol/g), indicating superior preservation against oxidative damage. In contrast, oranges packaged in the control and LDPE films showed higher MDA levels of 7.67 ± 0.29 nmol/g and 9.32 ± 0.53 nmol/g, respectively. The highest MDA content observed in oranges without packaging (17.82 ± 0.19 nmol/g), confirms significant oxidative stress and inferior fruit quality due to higher overripening (Table 5 & Fig. 12F).

#### 3.14.5. Fruit’s Antioxidant properties

Initially on Day 0 oranges inherently demonstrated antioxidant properties, with an initial radical scavenging activity (RSA) of 21.76 ± 2.86%. However, over a period of 7 days, this value declined to 11.01 ± 1.33%, which may be due to the continuous reaction between the endogenous antioxidants and the newly generated free radicals, thereby reducing its overall antioxidant capacity (Foroudi et al., 2014).

Oranges packaged in PGL film retained a significantly higher antioxidant activity of 22.17 ± 2.13%, suggesting that the radicals formed during storage were effectively neutralized by the PGL film, which possesses a strong antioxidant activity of 80.3 ± 0.43%. This also signifies that the PGL films can adsorb and quench free radicals through direct contact without releasing film components (lignin-associated antioxidant molecules). In contrast, oranges wrapped in the control film exhibited a reduced RSA of 14.51 ± 2.67%; however, this value is notably higher than that of oranges stored in LDPE film. The antioxidant capacity of oranges packaged in LDPE declined markedly to 10.99 ± 2.25%, underscoring the film’s ineffectiveness in maintaining the oxidative stability and overall quality of the fruit during storage (Table 5 & Fig. 12F).

#### 3.14.6. Sucrose content

Initially, on day 0, the sucrose content in fresh oranges was 3.6 ± 0.08% (w/w). After 7 days, a substantial decline was observed in oranges without packaging, with sucrose levels dropping to 1.28 ± 1.18% (w/w). This reduction can be attributed to increased respiration and microbial activity.

Oranges packaged in PGL films retained a sucrose content of 3.51 ± 0.67% (w/w), suggesting effective preservation attributed to the antimicrobial efficacy of the PGL film. This likely inhibited microbial proliferation and minimized respiratory activity, thereby reducing the metabolic consumption of sucrose. In comparison, oranges wrapped in control films showed a slightly decreased sucrose content of 3.13 ± 0.98% (w/w), indicating relatively lesser protection against sucrose degradation during storage. Meanwhile, oranges stored in commercial LDPE films exhibited a more substantial decline in sucrose content, reaching 2.83 ± 0.93% (w/w), which indicates that LDPE films were comparatively less effective in preserving sugar levels than both PGL and control films. (Table 5 & Fig. 12F).

#### 3.14.7. Ethylene concentration

Ethylene, a plant hormone, plays a central role in fruit overripening with its production rising significantly during the ripening process (Pretel et al., 1999). Ripening reduces fruit shelf life by altering parameters such as appearance, firmness, pH, malondialdehyde (MDA) content, and sucrose levels. Based on the above results, oranges without packaging exhibited the poorest quality, likely due to accelerated overripening, which corresponds to the higher ethylene emission rate observed per hour as shown in Fig. 12C.

After a 7-day storage period, oranges without packaging exhibited a marked increase in ethylene production, releasing 0.11 ± 0.004 ppm/g after 1 h, which further escalated to 0.14 ± 0.007 ppm/g after 7 h. In contrast, oranges enclosed in PGL films produced 0.03 ± 0.003 ppm/g of ethylene after 1 h which increased to 0.05 ± 0.003 ppm/g after 7 h, indicating a notable suppression of the ripening process. In oranges wrapped in control films, 0.07 ± 0.003 ppm/g of ethylene was produced after 1 h which rose to 0.11 ± 0.003 ppm/g after 7 h. Collectively, these observations, as presented in Table 5 and Fig. 12E, provide strong evidence that PGL films can effectively retard ethylene evolution and thereby delay fruit ripening.

## Conclusions

Sustainable food packaging utilizing an eco-friendly fabrication process to minimize environmental impact was developed. Lignin was extracted from cotton stalks using the acid-alkali method, with a purity of 93.93 ± 5.29% and yield of 48.22 ± 2.45% (w/w). This extracted lignin was then incorporated into a PDA/gelatin matrix to fabricate PGL biocomposite films, cured at 60°C, adhering to all green chemistry principles. The high network strength of the PGL films, arising from the synergistic effect of both physical and chemical crosslinking, was confirmed through rheological studies. This enhanced structural integrity translated into a notable tensile strength of 8.5 MPa and an elongation of 72%, indicating robust mechanical performance. Additionally, the films exhibited excellent water stability and moisture absorption characteristics to manage accidental water exposure or spillage. The PGL films also exhibited excellent UV-blocking properties (SPF 62.08; 98.39% UV-B blocking), along with strong antioxidant and antibacterial activities. Notably, application studies using Kinnow oranges revealed that PGL-wrapped oranges maintained superior physicochemical quality over seven days of storage compared to control and LDPE-packaged fruits. Collectively, these findings establish the potential of lignin-based, bioactive PGL films as sustainable, multifunctional packaging materials for extending the shelf life and quality of perishable citrus fruits during storage and transport.

## Supporting information

https://docs.google.com/document/d/1IyRujWiFrZB1uozRaewcxoPuugRWGSe1/edit?usp=sharing&ouid=105691328661985019790&rtpof=true&sd=true

## CRediT authorship contribution statement

**Ayan Banerjee:** Writing - original draft, Investigation, Methodology, Data curation, Validation, Conceptualization.

**Mohit Kumar Mehra**: Writing - original draft, Investigation, Data curation, Conceptualization.

**Althuri Avanthi**: Writing – review & editing, Validation, Supervision, Conceptualization Project administration, Funding acquisition.

## Declaration of competing interest

The authors declare that they have no known competing financial interests or personal relationships that could have appeared to influence the work reported in this paper.

## Acknowledgements

The authors gratefully acknowledge the Science and Engineering Research Board (SERB) (SRG/2023/000913) for funding this project. The authors also thank the Director of the Indian Institutes of Technology (IIT), Hyderabad, for providing necessary infrastructural facilities. MKM acknowledges the University Grants Commission (UGC), India, for their financial assistance in the form of the UGC-NET Junior Research fellowship (NTA Ref. No.: 211610107803)

## Appendix A. Supplementary data

### Data availability

Data will be made available on request.

---

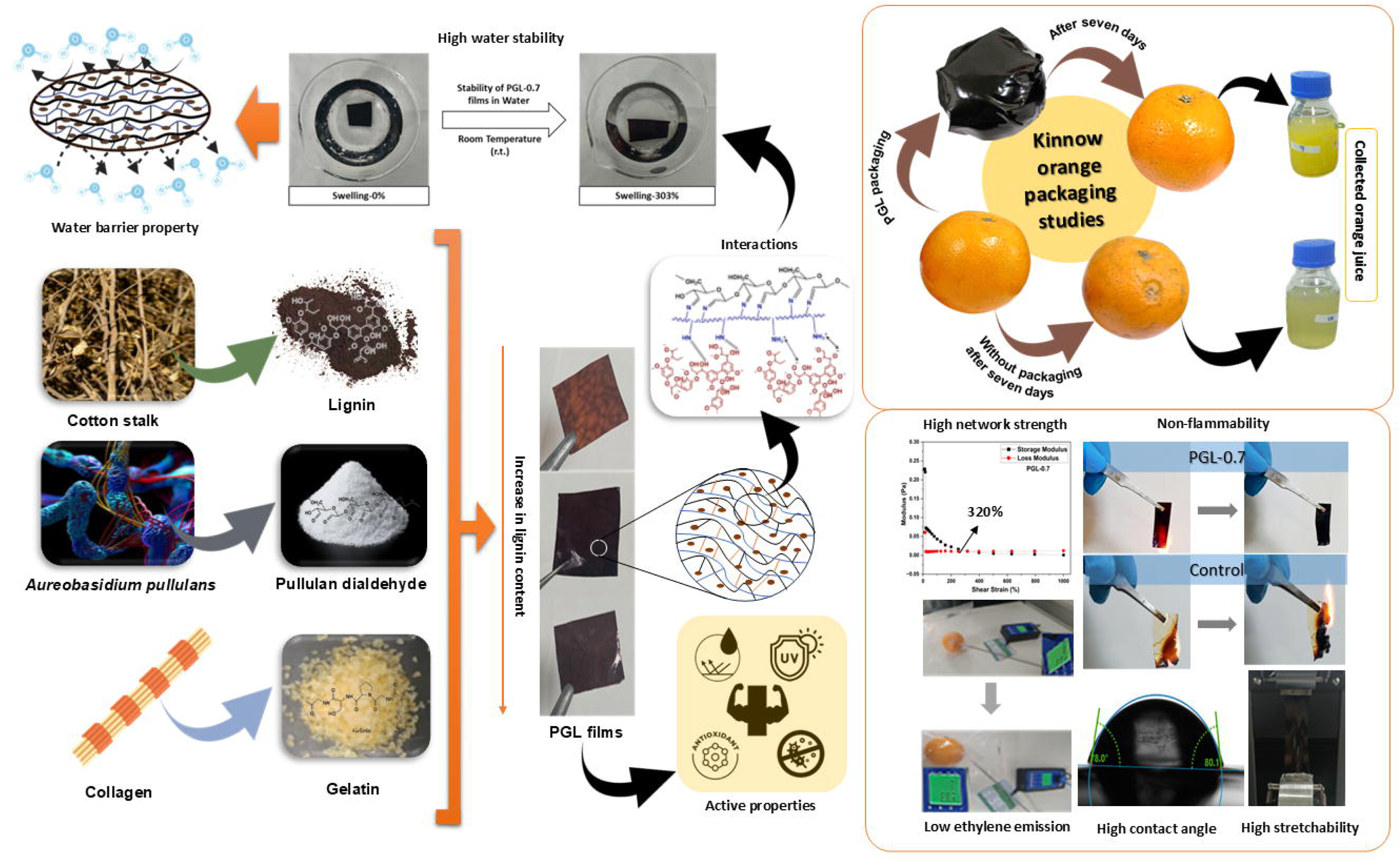

## Notes

### Competing Interest Statement

An Indian patent entitled - A bioactive green composite film and a method of preparation thereof, related to this biocomposite film formulation has been filed ((Application number 202541021083, Dated: 08-03-2025) and published (21-03-2025).

