## Supplementary material for "Green Synthesis of a Novel Gelatin Crosslinked Oxidized Pullulan-Lignin biocomposite film for active food packaging": https://docs.google.com/document/d/1IyRujWiFrZB1uozRaewcxoPuugRWGSe1/edit?usp=sharing&ouid=105691328661985019790&rtpof=true&sd=true

**Table S1 Compositional analysis of cotton stalk**

| **Components** | **Composition (%, w/w)** |
| --- | --- |
| Lignin | 25.44 ± 0.49 |
| Cellulose | 46.67 $\pm$ 0.49 |
| Ethanol extractives | 2.23 ± 0.82 |
| Ash | 4.09 ± 0.09 |

**List of figures**

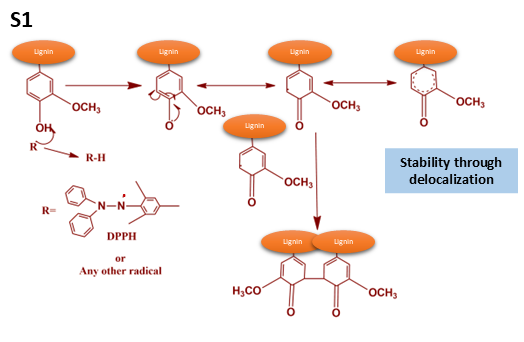

**Fig. S1** Free radical scavenging mechanism of Lignin

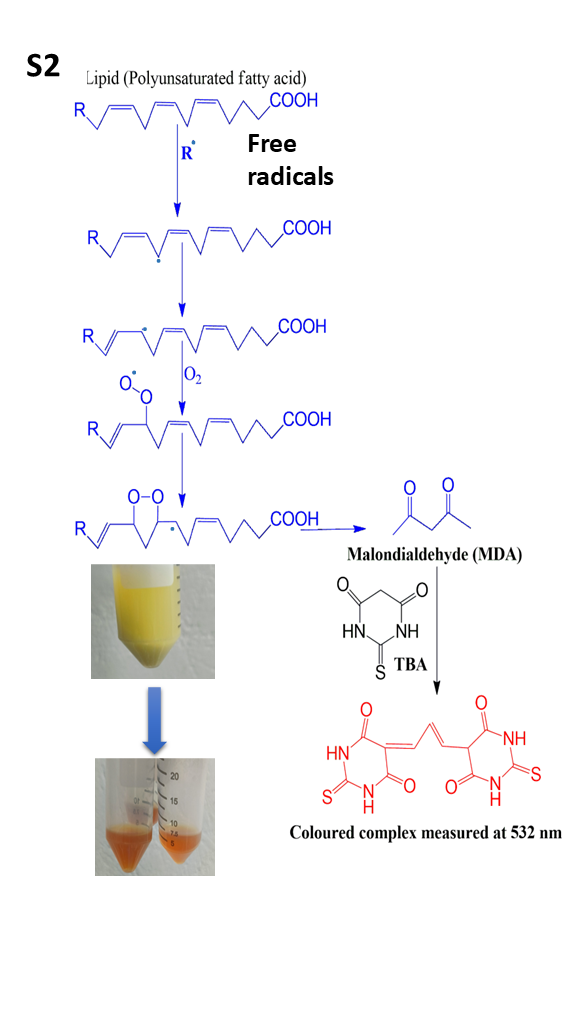

**Fig. S2** MDA formation scheme by lipid peroxidation
